# Glia-derived exosomal miR-274 targets Sprouty in trachea and synaptic boutons to modulate growth and responses to hypoxia

**DOI:** 10.1101/547554

**Authors:** Yi-Wei Tsai, Hsin-Ho Sung, Jian-Chiuan Li, Chun-Yen Yeh, Pei-Yi Chen, Ying-Ju Cheng, Chun-Hong Chen, Yu-Chen Tsai, Cheng-Ting Chien

**Affiliations:** Institute of Molecular Biology, Academia Sinica, Taipei, Taiwan; National Institute of Infectious Diseases and Vaccinology, National Health Research Institutes, Zhunan, Taiwan; Taiwan International Graduate Program in Interdisciplinary Neuroscience, National Cheng Kung University and Academia Sinica, Taipei, Taiwan; Institute of Neuroscience, National Yang-Ming University, Taipei, Taiwan; Department of Life Science, Tunghai University, Taichung, Taiwan; Neuroscience Program of Academia Sinica, Taipei, Taiwan

**Author notes:** Address correspondence to: Cheng-Ting Chien, Ph.D. Neuroscience Program of Academia Sinica, 128 Academia Road, Section 2, Nankang, Taipei 11529, Taiwan, 886-2-2787-3284.

**Keywords:** microRNA, glia, exosome, ESCRT, Rab11, Syx1A, development, hypoxia, Drosophila

## Abstract

Secreted exosomal miRNAs mediate inter-organ/tissue communication by downregulating gene expression, thereby modulating developmental and physiological functions. However, the source, route, and function have not been formally established for specific miRNAs. Here, we show that glial miR-274 non-cell autonomously modulates the growth of synaptic boutons and tracheal branches. Whereas precursor miR-274 was expressed in glia, mature miR-274 was secreted. miR-274 secretion to circulating hemolymph was detected in exosomes, a process requiring ESCRT components in exosome biogenesis and Rab11 and Syx1A in exosome release. miR-274 downregulated Sprouty to activate MAPK in synaptic boutons and tracheal branches, thereby promoting their growth. Expression of miR-274 solely in glia of a *mir-274* null mutant reset normal levels of Sprouty and MAPK, and hemolymphatic exosomal miR-274. *mir-274* mutant larvae were hypersensitive to hypoxia, which was suppressed by increasing tracheal branches. Thus, glia-derived miR-274 coordinates growth of synaptic boutons and tracheal branches to modulate larval hypoxia responses.

## Introduction

Cells communicate at multiple levels during development, from short-to long-range, between the same or different types of cells, and between different tissues/organs in the body. Long-range communication requires transport of signals, leading to coordinated growth and differentiation in multicellular organisms. Several mechanisms for transporting long-range signals from source to target have been identified, including transport by extracellular vesicles (EVs) derived from ligand-expressing cells (1, 2). These EVs originate from at least two sources; direct shedding of plasma membranes to form microvesicles, and secretion of intraluminal vesicles (exosomes) from multivesicular bodies (MVBs). Exosomal transportation has been better characterized due to the consistent size of the vesicles (30-100 nm in diameter), easy detection in the circulatory system, and well-characterized cargoes (3). Furthermore, the physiological functions and diseases associated with secreted exosomes have been studied in greater detail (4).

Secreted exosomes host a major type of cargo, i.e., non-coding microRNAs (miRNAs), which can functionally inhibit protein expression in target cells (3). In animals, miRNAs are small RNAs of ~22 nucleotides, which possess a seed region of typically 2-7 nucleotides at their 5’ ends that binds to sequences of the target mRNAs to promote mRNA degradation or translational repression (5). Although cell-autonomous functions of miRNAs have been amply reported, non-cell-autonomous functions have only been recently discovered. Circulating miRNAs can target gene expression in distant tissues. For example, during the formation of immune synapses, exosomal miR-335 is transferred from T cells to antigen-presenting cells to downregulate *SOX-4* mRNA translation (6). Exosomal miR-451 and miR-21 are transferred from glioblastoma to microglia to downregulate c-Myc expression (7). Adipocyte-derived exosomal miR-99b targets *Fgf21* in hepatic cells to downregulate mRNA and protein expression (8). In *Drosophila*, epithelial cells express *bantam* miRNA to regulate neuronal growth non-cell-autonomously (9). miRNAs have also been isolated from the circulating hemolymph of *Drosophila* (10), suggesting that some *Drosophila* miRNAs are carried by exosomes in the circulatory system to function systematically or in specific target cells. However, mechanistic details—such as the sources of exosomal miRNAs, their presence in circulating hemolymph, and their direct target genes in recipient tissues, as well as functional modulation of recipient tissues and relevant physiological functions—have not been established for a specific miRNA, especially in a model organism that would greatly facilitate a precise mechanistic understanding at the genetic level.

During vertebrate development, nerve and blood vessel formation share many cellular processes, including cone-like growth tips, branching patterns and ramifying networks (11, 12). Pairs of signals and receptors such as Slit and Robo, Netrin and Unc5/DCC co-receptor, and Ephrin and Eph, which were initially identified as being involved in axon outgrowth and patterning, have since been shown to be involved in similar processes in vasculogenesis (11, 12). Expression of vascular endothelial growth factor (VEGF), which plays critical roles in angioblast migration and vessel ingression, is spatiotemporally regulated in the neural tube during embryonic development (13). Although VEGF167 and the axon guidance signal Sema3A function separately in early vessel and nerve formation, both signals function through the shared receptor neurophilin-1 (14). During post-developmental stages, neuronal activity of the nervous system and oxygen delivery are also prominently coupled, forming the neurovascular units (15). Given the extreme sensitivity of the nervous system to alterations of ions, nutrients and potentially harmful molecules in the vascular system, an interface between both systems is necessary. Astrocytes, the major type of glia in the mammalian brain are structurally and functionally coupled to neuronal synapses and vascular endothelial cells to directly regulate their activities and communication (16–20). The insect trachea, the prototypical vascular system, allows oxygen delivery to the inner parts of the animal body. Nerves, glial sheath, and tracheal branches have been described for the larval brains and adult NMJs of *Drosophila* (21–23). Synapse organization and activity of larval NMJs, as well as their glial interactions, have also been well characterized (23–25).

In this study, we explored larval *Drosophila* neuromuscular junctions (NMJs) where axonal terminals branch out to form synaptic boutons with muscle membranes. We show that tracheal terminal branches are also highly arborized at the NMJs, making them an ideal system for studying coordinated nervous and vascular development. We screened a collection of miRNA knockout mutants and identified the *mir-274* mutant as having defects in both synaptic and tracheal growth. By fluorescent *in situ* hybridization (FISH), we showed that the miR-274 precursor was expressed in glia and the mature form was ubiquitously detected. Consistently, miR-274 was required in glia for synaptic and tracheal growth. Glial expression of miR-274 could be detected in the hemolymph of the larval circulatory system. Indeed, miR-274 was secreted as an exosomal cargo as shown by genetic analysis and biochemical fractionation. We identified a miR-274 target *sprouty* (*sty)* that includes a targeting site in the 3’UTR of the *sty* transcript and showed that glial miR-274 expression induced Sty downregulation and MAPK activation in synaptic boutons and tracheal branches. Intriguingly, the *mir-274* mutant that had fewer tracheal branches were hypersensitive to hypoxia, and both phenotypes were suppressed *sty* loss-of-function mutations. Thus, we demonstrate the non-cell-autonomous developmental role of glial miR-274, which coordinates the growth of synaptic boutons and tracheal branches and is required for normal responses to hypoxia.

## Results

### miR-274 is required in glia to modulate synapse and tracheal growth

By immunostaining synaptic, glial and tracheal structures, we show that glial processes wrap around incoming motor axons, and envelopment ends before terminal branching at muscle 6/7 (Figures S1A & A’, arrowheads). Axonal terminal branches form bouton-like structures that innervate muscles to form functional synapses, and multiple tracheal branches terminate near these synaptic boutons (Figure S1A”, empty arrowheads). Ultrastructure analysis by transmission electron microscopy shows that glial processes enwrap axonal processes close to tracheal branches (Figure S1B, arrows). Synaptic boutons wrapped in the subsynaptic reticulum (SSR) are also visible (Figure S1B, arrowhead). These observations suggest that the glia-synapse-trachea organization might represent functional units and its formation might be developmentally regulated.

To investigate whether these structures are developmentally co-regulated, we screened a collection of miRNA knockout mutants (26). We identified *mir-274*^*KO*^ as displaying reduced growth of both synaptic boutons and tracheal branches (Figure 1A). Quantification revealed that larvae homozygous for *mir-274*^*KO*^ exhibited a ~40% reduction in the numbers of synaptic boutons and tracheal terminal branches compared to both wild-type *w*^*1118*^ and *mir-274*^*KO*^/+ larvae (Figure 1D). We also detected reduced numbers of synaptic boutons and tracheal branches for the trans-heterozygous *mir-274*^*KO*^/*mir-274*^*6-3*^ mutant (Figures 1A and 1D), confirming that the lack of miR-274 activity accounts for growth defects in both systems. This reduction in synaptic bouton number was not limited to NMJs of muscle 6/7, as we also observed bouton reduction at NMJs of muscle 4 in the *mir-274*^*KO*^ mutant (Figures S1C and S1D). Likewise, tracheal branching was also reduced in the dorsal region of the *mir-274*^*KO*^ mutant (Figures S1C and S1D). These data suggest that larvae lacking miR-274 fail to develop complete sets of synaptic boutons and tracheal branches.

**Figure 1.**
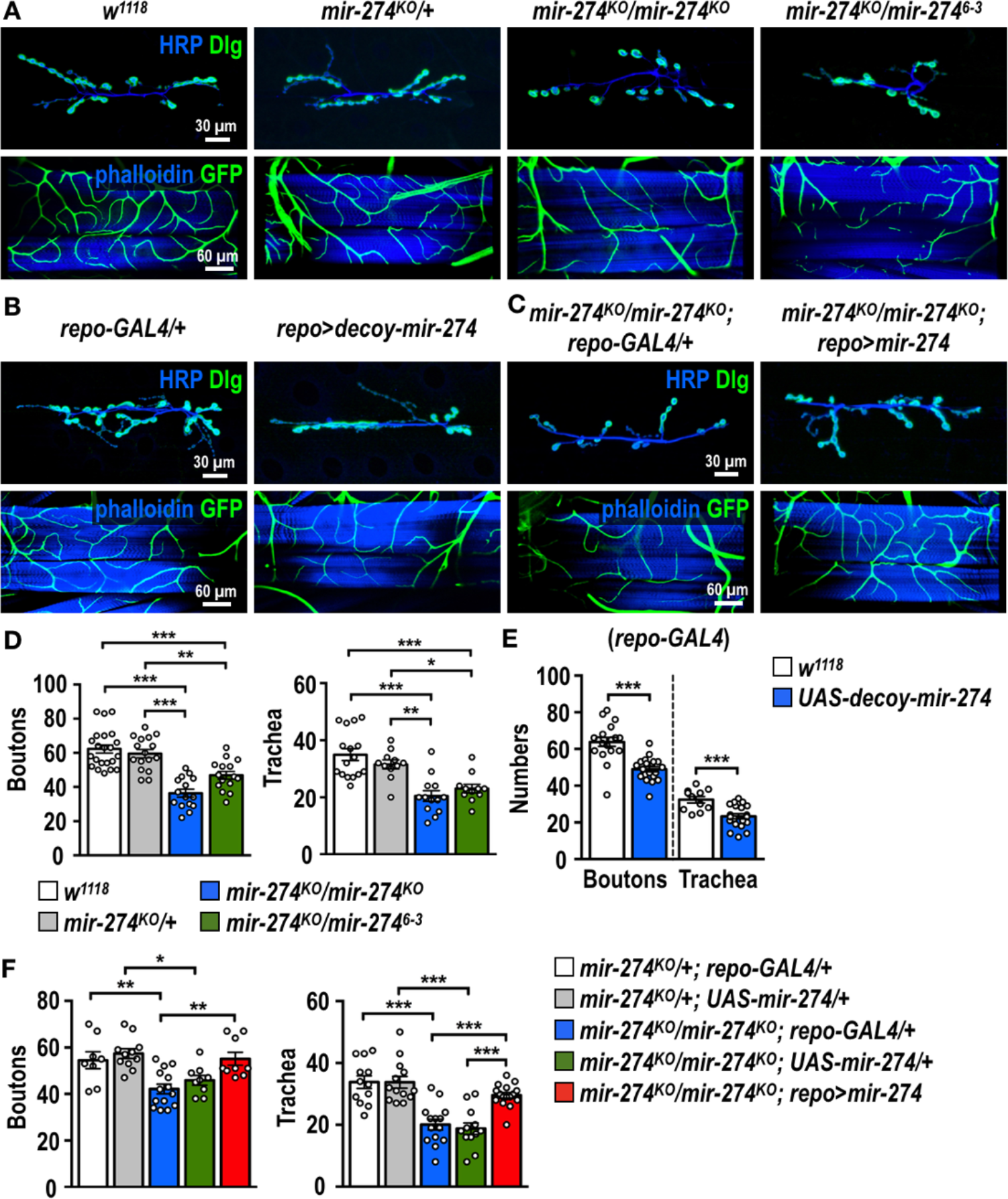
Reduced synaptic boutons and tracheal branches in miR-274 mutants. (A-C) Images of synaptic boutons (top rows) immunostained for presynaptic HRP (blue) and postsynaptic Dlg (green), and tracheal branches (bottom rows) revealed by *btl-lexA*>*lexAop-CD2-GFP* (green) counterstained for muscle phalloidin (blue), for (A) *w*^*1118*^ and *mir-274*^*KO*^/+, *mir-274*^*KO*^/*mir-274*^*KO*^, and *mir-274*^*KO*^/*mir-274*^*6-3*^, (B) *repo-GAL4* control (crossed to *w*^*1118*^, left panels), and *repo-GAL4* crossed to *UAS-decoy-mir-274* (*repo*>*decoy-mir274*, right panels) for miR-274 depletion, and (C) the miR-274 mutant (*mir-274*^*KO*^/*mir-274*^*KO*^; *repo-GAL4*/+) and glial rescue (*mir-274*^*KO*^/*mir-274*^*KO*^; *repo*>*mir-274*). Scale bars are shown at bottom right, with 30 μm for synaptic bouton images and 60 μm for tracheal branch images. (D-F) Bar graphs showing mean ± SEM (with dots representing data points) to compare numbers of synaptic boutons and tracheal branches for genotypes in (A-C). Statistical significance was assessed by One-way ANOVA followed by Tukey post hoc tests for (D, F), or Independent *t*-tests (E), and shown as * for *p* < 0.05, ** for *p* < 0.01, and *** for *p* < 0.001.

To examine whether specific types of cells require miR-274 for growth of synaptic boutons and tracheal branches, we employed the *UAS-decoy-mir-274* transgene (see Experimental Procedures) driven by cell-type specific *GAL4* drivers to inhibit miR-274 functions. Neuronal *elav-GAL4*, glial *repo-GAL4*, and tracheal *btl-GAL4* were individually crossed to *UAS-decoy-mir-274* to analyze synaptic bouton and tracheal branch phenotypes. Surprisingly, although miR-274 inhibition by neuronal and tracheal drivers had no obvious phenotypic impact (Figures S1E-H), glial depletion caused significant reductions in synaptic bouton and tracheal branch numbers (Figures 1B and 1E). Whereas the glial processes at NMJs of muscle 6/7 presented a normal morphology in the *mir-274*^*KO*^ mutant, the hemolymph brain barrier (HBB, which is mainly composed of glia) was defective (Figure S1I), confirming the findings of a previous study (26). However, the reductions in synaptic boutons and tracheal branches are not secondary to the defective HBB, as an intact HBB was retained in glial expression of *UAS-decoy-mir-274* (Figure S1I) while synaptic boutons and tracheal branches were reduced (Figures 1B and 1E). Thus, it seems that glial inhibition of miR-274 is sufficient to compromise synapse and tracheal growth.

Furthermore, we performed a glial rescue experiment. In the homozygous *mir-274*^*KO*^ mutant carrying either *repo-GAL4* or *UAS-mir-274* alone, the numbers of synaptic boutons and tracheal branches were fewer than the numbers in the heterozygous mutants carrying either *repo-GAL4* or *UAS-mir-274* alone (Figures 1C and 1F). However, in the homozygous *mir-274*^*KO*^ mutant carrying both *repo-GAL4* or *UAS-mir-274*, the numbers of synaptic boutons and tracheal branches were comparable to those in the heterozygous mutants (Figures 1C and 1F). These results strongly support that glia-expressed miR-274 promotes growth of synaptic boutons and tracheal branches.

### Expression of miR-274 precursor and mature forms

To characterize miR-274 expression, we performed FISH experiments using probes complementary to the loop or the stem sequence to detect the precursor or the mature forms of miR-274 (Figure 2A), respectively, in dissected larval fillets (Figure 2B). In control, we only detected low background or non-specific signals in larval brain using scrambled probes (Figure 2C). However, the loop probe for detecting the miR-274 precursor presented prominent signals in the brain (Figures 2D and 2D’). These signals were localized in glia labeled with *repo-GAL4*-driven *mCD8-GFP* and occasionally strong nuclear signals were detected (Figure 2D’, arrowhead). In contrast, low background signals were detected in synaptic boutons and tracheal branches (Figures 2F, 2F’, 2H and 2H’). We then employed the stem probe to detect mature miR-274 (Figure 2A). Interestingly, we observed strong and ubiquitous signals, i.e., not restricted to specific cells in the brain (Figure 2E and 2E’). These signals were also detected in muscle cells and within synaptic boutons (Figures 2G and 2G’), as well as in tracheal soma and branches (Figures 2I and 2I’). Signals of mature miR-274 were not detected in the *mir-274*^*KO*^ mutant control (Figure 3G). Taken together, these results are consistent with the idea that the miR-274 precursor is mainly synthesized in glia and that the mature form is detected in muscles, synaptic boutons and tracheal cells.

**Figure 2.**
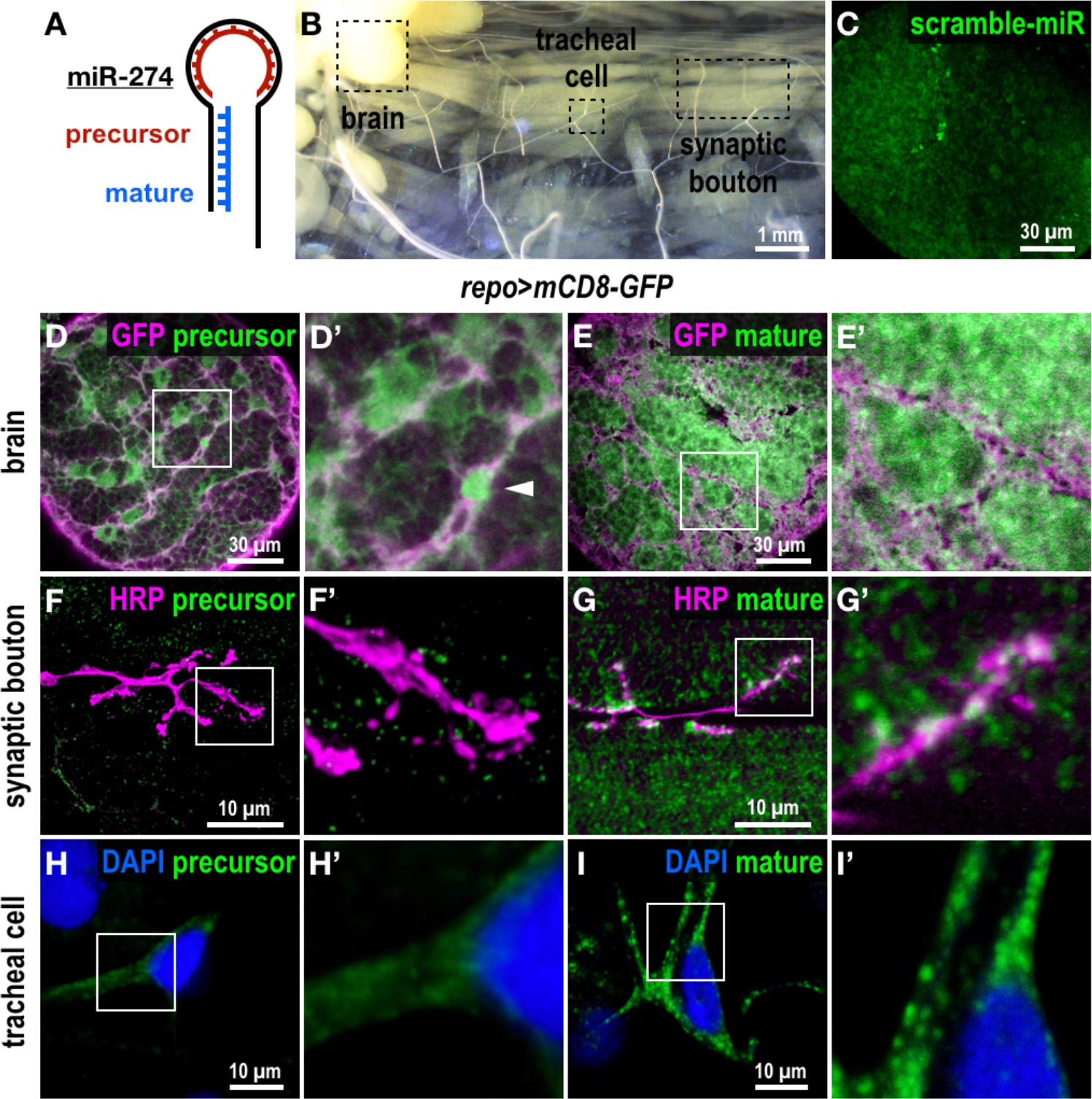
Expression of precursor and mature miR-274. (A) Illustration of the two FISH probes used to detect miR-274 expression. The precursor (pre-) probe compliments the loop region, and the mature probe compliments the stem miR-274 sequence. (B) Dissected third instar larvae (*w*^*1118*^) showing sites for detecting FISH signals in brain lobes, synaptic boutons and tracheal cells. (C-I) Images for FISH signals using the scramble (C), precursor (D, F and H) or mature probe (E, G and I), with glia labeled by *repo*>*mCD8-GFP* (magenta in D and E), synaptic boutons by HRP (magenta in F, G), and DAPI-labeled nuclei (blue in H, I). (C) Low background levels of non-specific FISH signals detected by the scramble probe in the brain lobe. (D) The precursor probe detected a glial pattern overlapping glial membrane (enlarged image in D’) with a glial nucleus indicated by an arrowhead. (F, H) Only diffusive or low background signals were detected within synaptic boutons (F and F’) and tracheal cells (H and H’). (E, G and I) The mature probe detected ubiquitous punctate signals in brain (E and E’), synaptic boutons and muscles (G and G’), and tracheal soma and branches (I and I’).

**Figure 3.**
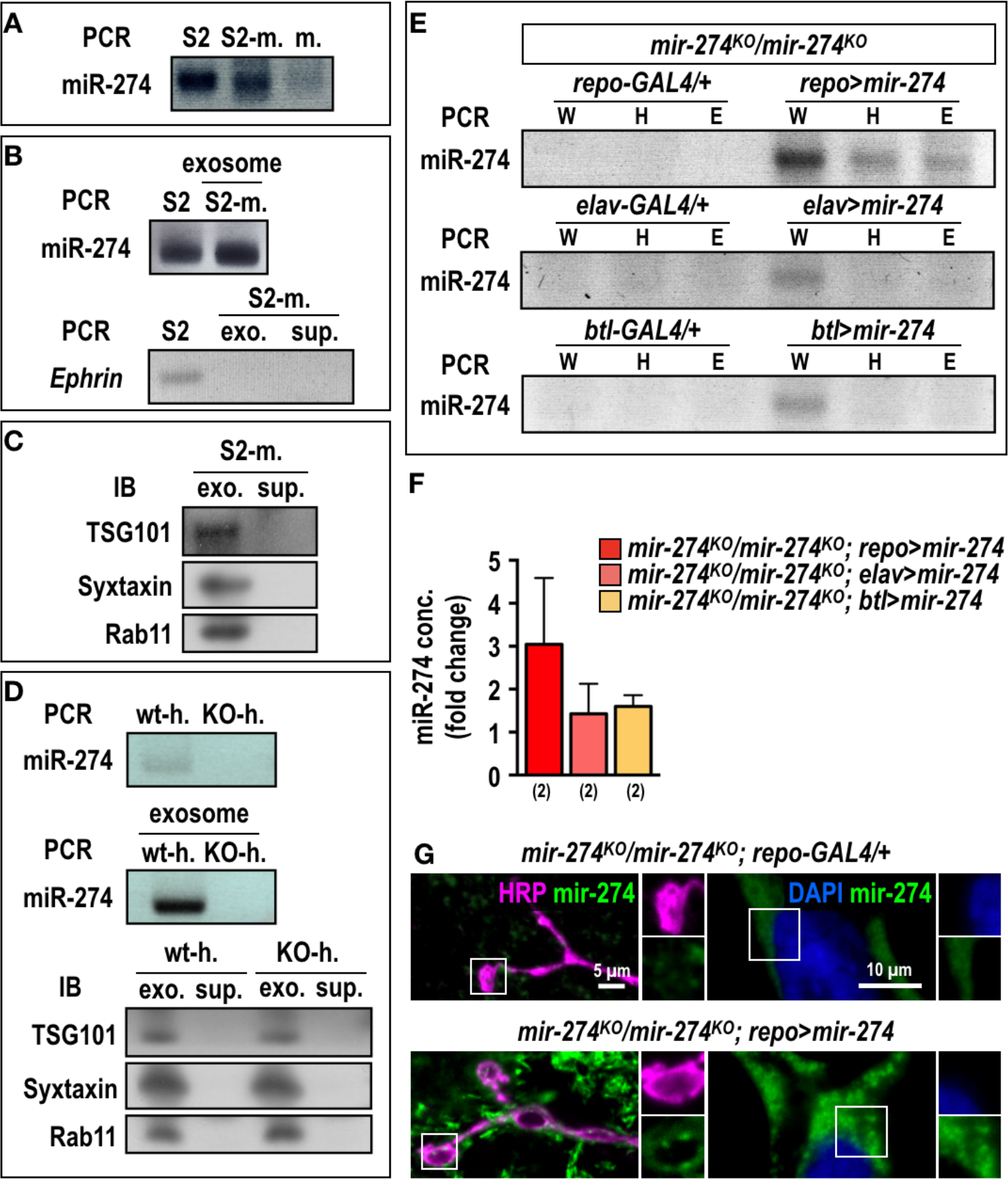
Secretion of exosomal miR-274 from glia. (A) miR-274 was detected by RT-PCR in S2 cells (S2) and medium used to culture S2 cells (S2-m.), but no signal was detected in fresh medium (m.). (B) S2 cells and isolated exosomal fractions from the culture medium also express miR-274 as detected by RT-PCR. *Ephrin* transcript is detected in whole S2 cell lysates and absent in exosomal fractions (exo.) and supernatant (sup.) from cultured S2 medium (B). Immunoblots showing the presence of TSG101, Syxtaxin, and Rab11 in exosomal fractions isolated from cultured S2 medium but not in supernatants (C). (D) miR-274 was detected in *w*^*1118*^ hemolymph (h.), and its exosomal fractions, but was not detected in *mir-274*^*KO*^ hemolymph and the exosomal fractions. Presence of TSG101, Syxtaxin and Rab11 in exosomal fractions isolated from *w*^*1118*^- and *mir-274*^*KO*^-hemolymph, but not in supernatants. (E) Detection of miR-274 in the *mir-274*^*KO*^ mutant with or without expression of *UAS-mir-274* by *repo-GAL4* (top), *elav-GAL4* (middle) or *btl-GAL4* (bottom) in whole larval lysates (w), hemolymphs (h) and exosomal fractions (e). (F) Quantification of miR-274 levels by absolute qPCR in larval exosomal fractions in the *mir-274*^*KO*^ mutant hosting different *GAL4*-driven miR-274 expression, which were normalized to the *mir-274*^*KO*^ mutant carrying the respective *GAL4* driver alone. (G) Images show background miR-274 FISH signals by the mature probe in mutant (*mir-274*^*KO*^/*mir-274*^*KO*^; *repo-GAL4*) and brighter punctate signals in glial rescue (*mir-274*^*KO*^/*mir-274*^*KO*^; *repo*>*mir-274*) larvae.

### Exosomal secretion of miR-274 from glia

To examine how miR-274 is secreted, we first examined whether miR-274 could be secreted from S2 cells. Indeed, we detected miR-274 in S2 cell extracts (Figure 3A). Significant levels of miR-274 could also be detected in the medium used to culture S2 cells, but not in the medium in which S2 cells were not cultured, indicating that miR-274 could be secreted from S2 cells into the medium (Figure 3A). To examine whether secreted miRNA is through secretory exosomes, we fractionated and pelleted the S2 cell culture medium to enrich for exosomes. Similar to whole cell extracts, the fractionated exosomes were enriched with miR-274 (Figure 3B). As a positive control, the exosomal markers TSG101, Rab11 and Syntaxin (27, 28) were detected in the exosomal fraction, and not detected in the exosome-depleted supernatant (Figure 3C). As negative controls, *Ephrin* mRNA that is not present in the secreted exosomes (29) was only detected in whole cell extracts and not in the exosome fraction or in the exosome-depleted supernatant (Figure 3B).

As miRNAs could be transported by circulating exosomes, we then examined whether miR-274 could be detected in the larval hemolymph. Significant amounts of miR-274 were detected in the hemolymph of wild-type larvae, which were absent in the hemolymph isolated from the *mir-274*^*KO*^ mutant (Figure 3D). We fractionated and pelleted exosomes from the hemolymph and found that fractionated exosomes were enriched with miR-274 (Figure 3D). The exosomal fraction isolated from the *mir-274*^*KO*^ hemolymph did not contain miR-274 (Figure 3D). The typical exosomal markers TSG101, Rab11 and Syntaxin were detected in exosomal fractions derived from both hemolymphs of wild-type control and mutant larvae but not in the exosome-depleted supernatants (Figure 3D). Thus, miR-274 could be secreted into larval hemolymph and S2 cell culture medium as circulating exosomes.

With the detection of miR-274 in the exosomes of hemolymphs, we would like to detect whether glia could secret miR-274 carried by exosomes to the hemolymph. As *mir-274*^*KO*^ is a deletion allele, no miR-274 could be detected in whole larval lysates, hemolymphs and hemolymph-derived exosomal fractions in the homozygous *mir-274*^*KO*^ mutant (*mir-274*^*KO*^/*mir-274*^*KO*^; *repo-GAL4*, Figure 3E). In contrast, miR-274 was detected in all three of these preparations from the homozygous *mir-274*^*KO*^ larvae carrying both *repo-GAL4* and *UAS-mir-274* (*mir-274*^*KO*^/*mir-274*^*KO*^; *repo*>*mir-274*, Figure 3E). Thus, glia could secrete miR-274 as an exosomal cargo in the hemolymph. We performed the same sets of experiments for neuronal and tracheal miR-274 expression in the *mir-274*^*KO*^ mutant. miR-274 was only detected in whole larval lysates, but not in the isolated hemolymphs or exosomal fractions (Figures 3E). Quantification of miR-274 levels by absolute qPCR (Experimental Procedures) in hemolymph-derived exosomal fractions showed a three-fold increase in *mir-274*^*KO*^/*mir-274*^*KO*^; *repo*>*mir-274* as compared to *mir-274*^*KO*^/*mir-274*^*KO*^; *repo-GAL4* whereas *elav-GAL4*- and *btl-GAL4*-driven expressions were comparable to respective *GAL4* driver controls (Figure 3F). Therefore, glia is the major source to release exosomal miR-274 into the hemolymph.

To further show that glial expression of miR-274 could reach synaptic boutons and tracheal branches for function, we performed the FISH experiment with the mature miR-274 probe. In the control of the *mir-274*^*KO*^ null mutant carrying only *repo-GAL4* (*mir-274*^*KO*^/*mir-274*^*KO*^; *repo-GAL4*), low background FISH signals appeared to be homogeneously throughout different tissues, including synaptic boutons, muscle cells, and tracheal cells (Figure 3G). This confirms the specificity of the mature probe in detecting miR-274. In glial expression of miR-274 in the *mir-274*^*KO*^ null mutant (*mir-274*^*KO*^/*mir-274*^*KO*^; *repo*>*mir-274*), we detected strong and punctate signals in synaptic boutons, muscle cells, and tracheal cells (Figure 3G). We also examined neuronal expression of *UAS-mir-274* in the *mir-274*^*KO*^ mutant, which showed strong miR-274 FISH signals in synaptic boutons (Figure S2A). However, only background signals were detected in tracheal cells (Figure S2A). Likewise, miR-274 FISH signals were detected only in tracheal cells and not in synaptic boutons of *mir-274*^*KO*^ mutants upon tracheal *UAS-mir-274* expression (Figure S2D). Taken together, these results suggest that miR-274 expressed in glia could be secreted and detected in the target sites like synaptic boutons and tracheal cells.

### Exosomal secretion of miR-274 from glia requires ESCRT components, Rab11, and Syx1A

Exosomal release requires Rab11 in MVB transportation and Syntaxin 1A (Syx1A) in membrane fusion with the plasma membrane (30). To show that miR-274-carrying exosomes are secreted through the process, we expressed *Rab11*^*RNAi*^ or *Syx1A*^*RNAi*^ in glia by *repo-GAL4*. We observed dramatically reduced miR-274 transcript levels in exosomes isolated from the hemolymph compared to the driver control (Figure 4A). To further confirm the absence of miR-274 by inhibiting glial exosomal secretion, we performed FISH with the mature miR-274 probe. In the *repo-GAL4* control, miR-274 signals delineated bouton morphology and were strongly punctate in muscle and tracheal cells (Figure 4B). However, these miR-274 signals were not detected upon glial knockdown of Rab11 or Syx1A (Figure 4B). The failure to detect miR-274 in the hemolymphs and target sites in Rab11 and Syx1A knockdowns is consistent with the idea that the exosomal release pathway is essential for miR-274-carried exosomes to be released from glia. We then examined whether the growth of synaptic boutons and tracheal branches, the target sites of miR-274, were affected in the inhibition of exosomal release. As expected, glial knockdowns of Rab11 or Syx1A caused reduced growth of synaptic boutons and tracheal branches (Figures 4C and 4D). Thus, just like the lack of miR-274 in glia, the lack of Rab11 or Syx1A in glia causes synapse and tracheal undergrowth. Cargo-carrying exosomes are assembled through serial actions of the ESCRT complexes, which promote membrane invagination and formation of intraluminal vesicles in MVBs (27). To confirm that miR-274 is packaged as exosomal cargo, we examined whether disruption of ESCRT complex components in glia could affect the level of circulating miR-274. We chose to knock down TSG101 of the ESCRT-I complex and Shrb of the ESCRT-III complex as RNAi knockdown of TSG101 or Shrb in glia resulted in efficient suppression of synapse and tracheal growth (Figures S3B and S3C). When we specifically expressed *TSG101*^*RNAi*^ or *shrb*^*RNAi*^ in glia by *repo-GAL4*, we observed reduced levels of miR-274 in exosomes isolated form the hemolymph, as compared to the driver control (Figure S3A). Thus, the glial ESCRT complex is also involved in the presence of miR-274-carrying exosomes in the hemolymph. Taken together, these results strongly suggest that miR-274 is secreted as an exosomal cargo to be released into the hemolymph to modulate synapse and tracheal growth.

**Figure 4.**
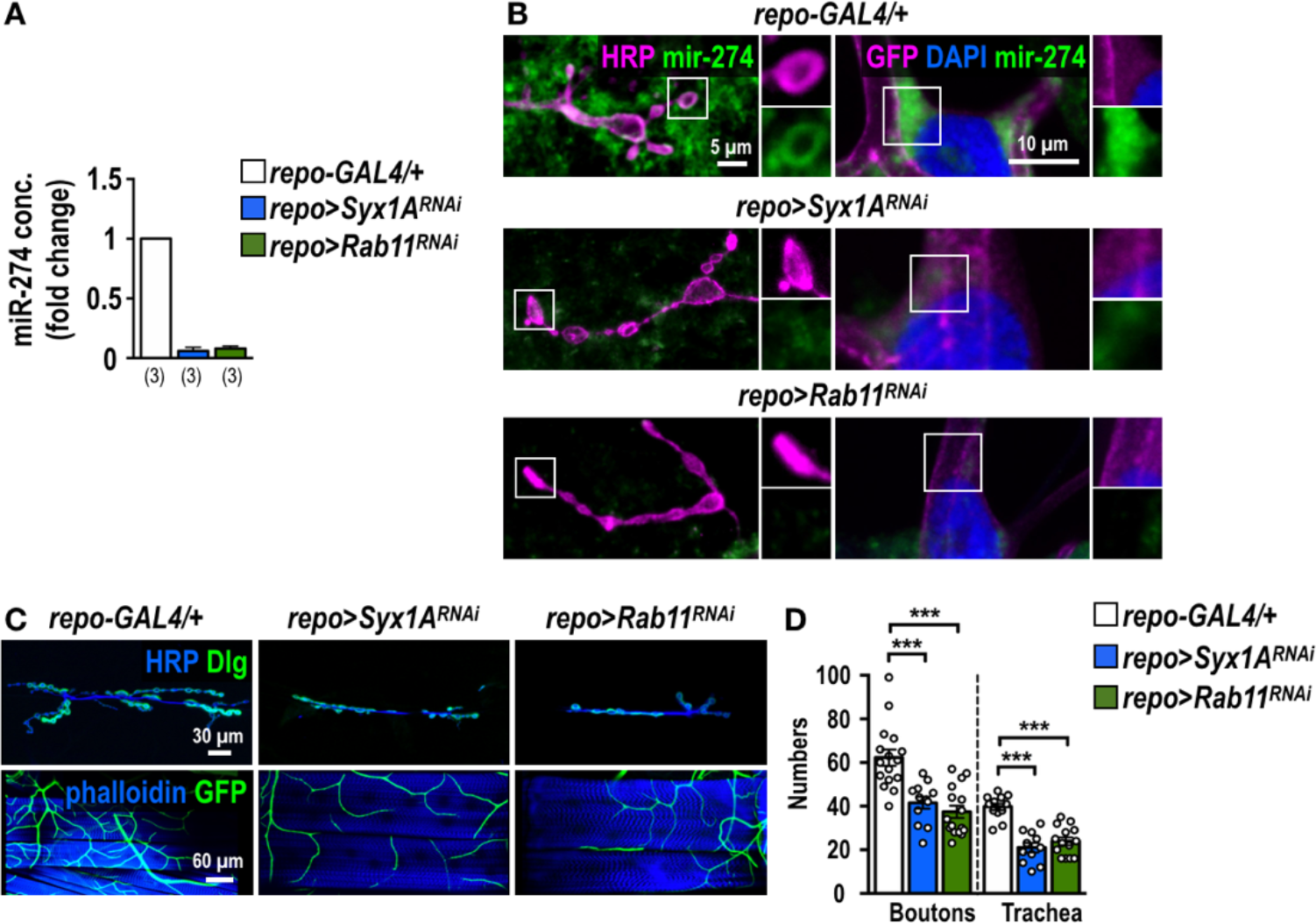
Secretion of exosomal miR-274 requires Rab11 and Syx1A in glia. (A) Absolute qPCR was performed to detect exosomal miR-274 in hemolymph prepared from *repo-GAL4* and knockdowns of Syx1A and Rab11. (B) Images show FISH signals for mature miR-274 in synaptic boutons and tracheal cells of *repo-GAL4* control (top row), which are largely lacking in *repo*>*Syx1A*^*RNAi*^ (middle) and *repo*>*Rab11*^*RNAi*^ (bottom). (C) Confocal images of synaptic boutons and tracheal branches in *repo-GAL4* control, *repo*>*Syx1A*^*RNAi*^ and *repo*>*Rab11*^*RNAi*^. (D) Bar graph for quantification of synaptic boutons and tracheal branches. Data were analyzed by One-way ANOVA followed by Tukey post hoc tests.

### *sprouty* as a target gene for miR-274 regulation

To understand how miR-274 regulates synapse and tracheal growth, we searched for genes that harbor miR-274 target sites and exhibited upregulated expression in the *mir-274*^*KO*^ larvae (Figure S4A). From among the resulting candidate genes (Figure S4B), we chose *sprouty* (*sty*) for further study since Sty plays a critical role in feedback inhibition of receptor tyrosine kinase (RTK) signaling during tracheal branching and synaptic bouton formation (31–34). The 3’UTR of *sty* mRNA contains a target site for miR-274 recognition. We then generated two luciferase reporter transgenes with either precise or mismatched miR-274 sequences targeting the *sty* 3’UTR (Figure 5A). As expected, the precise miR-274 targeting sequence downregulated reporter activity (relative to the vector control) when it was co-transfected with miR-274 (Figure 5A). The mismatched reporter was not downregulated upon miR-274 co-transfection. Thus, *sty* mRNA levels might be regulated by miR-274 through its 3’UTR targeting sequence. We then addressed whether miR-274 regulates *sty* mRNA expression *in vivo*. Indeed, higher *sty* transcript levels were detected in *mir-274*^*KO*^ larvae compared to the levels in wild-type control (Figure 5B), consistent with miR-274 having a role in downregulating *sty* expression.

**Figure 5.**
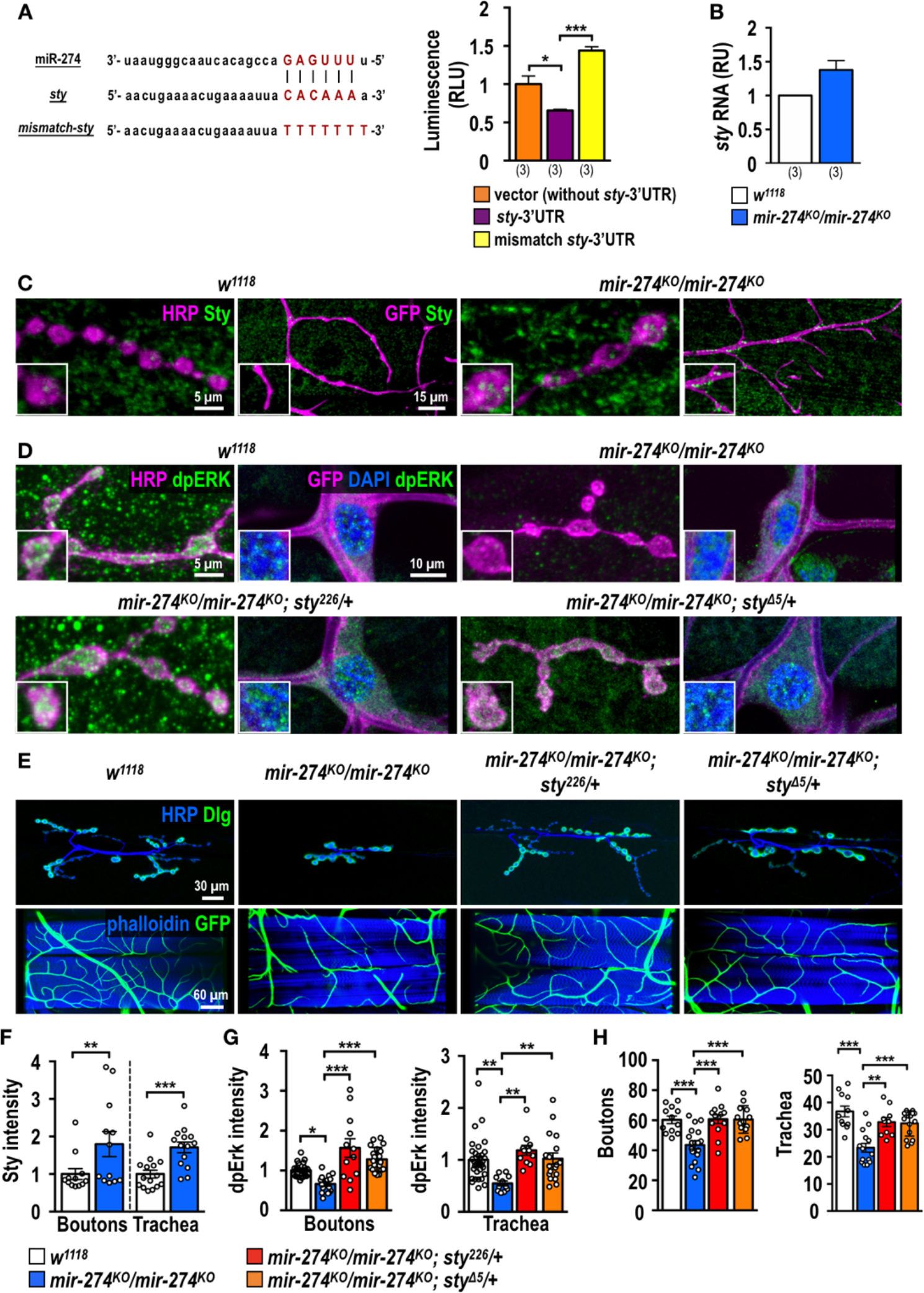
Regulation of Sty and dpERK expression by miR-274 in synaptic boutons and tracheal cells. (A) The miR-274 sequence with the seeding sequence in red, the predicted miR-274 targeting sequence at the *sty* 3’UTR with the complementary sequence in red, and the mismatch sequence with the mutated sequence in red. Luciferase reporter assay for vector control, vector with the targeting sequence, and vector with the mismatch sequence. Relative luciferase units (RLU) were normalized to an internal control. Data was analyzed by One-way ANOVA followed by Tukey post hoc tests. (B) The *sty* transcript levels in *w*^*1118*^ and *mir-274*^*KO*^ larval extracts were measured by qPCR and normalized to *Rpl19*, with the relative unit (RU) of *w*^*1118*^ set as 1. (C and D) Confocal images showing immunostaining of Sty (C) or dpERK (D) in synaptic boutons and tracheal cells in *w*^*1118*^, *mir-274*^*KO*^/*mir-274*^*KO*^ (C and D) and upon *sty* suppression in *mir-274*^*KO*^/*mir-274*^*KO*^; *sty*^*226*^/+ and *mir-274*^*KO*^/*mir-274*^*KO*^; *sty*^*Δ5*^/+ mutant larvae (D). (E) Images show synaptic boutons and tracheal branches in *w*^*1118*^, *mir-274*^*KO*^/*mir-274*^*KO*^, *mir-274*^*KO*^/*mir-274*^*KO*^; *sty*^*226*^/+, and *mir-274*^*KO*^/*mir-274*^*KO*^; *sty*^*Δ5*^/+. (F-H) Bar graphs for quantifications of Sty (F) and dpERK (G) immunofluorescence intensities within synaptic boutons and tracheal cells, and numbers of synaptic boutons and tracheal branches in *sty* suppression (H). (F) Data were analyzed by Independent *t*-tests. (G and H) Data were analyzed by One-way ANOVA followed by Tukey post hoc tests.

We further confirmed that Sty is regulated by miR-274 in synaptic boutons and tracheal branches by performing immunostaining. We first detected Sty expression in synaptic boutons and tracheal branches of the wild-type control. Levels of Sty in both these sites were enhanced relative to the control in the *mir-274*^*KO*^ mutant (Figure 5C), which is supported by quantifications of Sty immunofluorescence intensities (Figure 5F). Sty expression was also upregulated in muscle cells, suggesting that miR-274 might exert systemic regulation in multiple tissues (Figure 5C, also see Discussion). As a negative regulator, Sty inhibits several downstream components in RTK signaling, leading to downregulation of MAPK activity and inhibition of tissue growth (32–34). We further examined RTK/MAPK signaling activity by immunostaining for diphosphorylated-ERK (dpERK) (34). Levels of dpERK in synaptic boutons and tracheal cells of *mir-274*^*KO*^ were greatly reduced as compared to wild-type control (Figures 5D and 5G). Downregulation of dpERK levels is dependent on Sty, as elimination of one copy of *sty* in *mir-274*^*KO*^ (*mir-274*^*KO*^/*mir-274*^*KO*^; *sty*^*226*^/+ or *mir-274*^*KO*^/*mir-274*^*KO*^; *sty*^*Δ5*^/+) restored dpERK levels in tracheal cells and synaptic boutons to levels comparable to the control (Figures 5D and 5G). Restoration of dpERK levels was also detected in muscle (Figure 5D). We then tested whether miR-274 negatively regulates Sty expression to modulate synaptic and tracheal growth. Indeed, reducing the *sty* gene dosage in the miR-274 mutant suppressed both growth phenotypes (Figures 5E and 5H). These data support that miR-274 inhibits Sty expression, which leads to MAPK activation to promote the growth of synaptic boutons and tracheal branches.

### Glia-derived exosomal miR-274 targets Sty within synaptic boutons and trachea to modulate their growth

To understand whether glia-derived miR-274 regulates Sty and dpERK levels within synaptic boutons and tracheal cells, we first performed immunostaining in *repo*>*decoy-mir-274* larvae that had reduced synaptic and tracheal growth (Figures 1B and 1F). Similarly, trapping miR-274 in glia induced higher Sty levels in synaptic boutons and tracheal branches compared to *repo-GAL4* (Figures 6A and 6E). We also detected reduced levels of dpERK at these two sites (Figures 6B and 6F). In glia rescue larvae (*mir-274*^*KO*^/*mir-274*^*KO*^; *repo*>*mir-274*), as expected, we found reduced Sty and increased dpERK levels upon glial rescue (Figures 6C, 6D and 6G) and accompanied with increased numbers of synaptic boutons and tracheal branches (Figures 1C and 1F). These results strongly support that glia-expressed miR-274 reaches target sites to downregulate Sty expression and to promote growth of synaptic boutons and tracheal branches.

**Figure 6.**
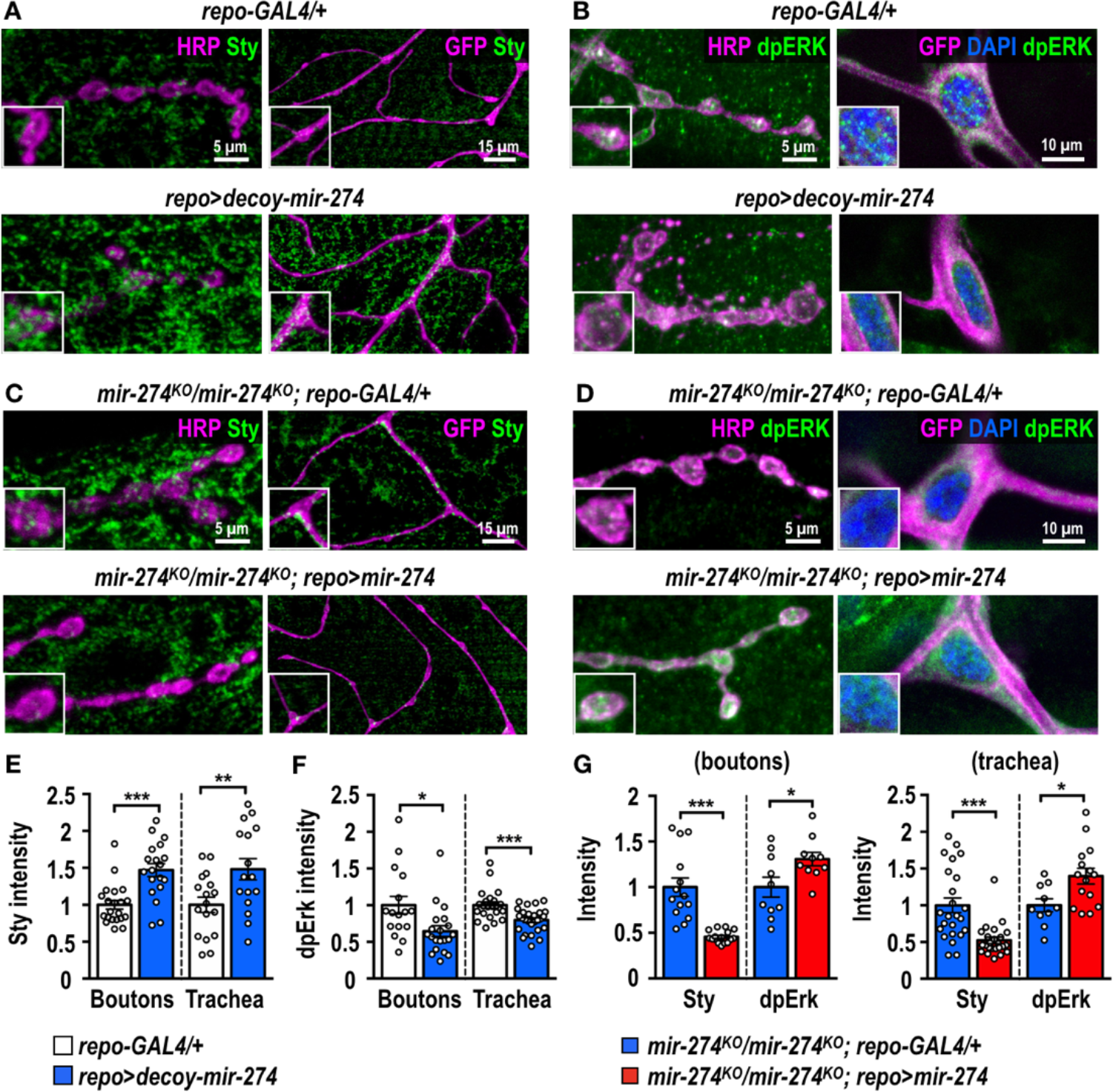
Glia-derived exosomal miR-274 modulates synaptic and tracheal growth. (A-D) Confocal images showing immunostaining of Sty (A and C) or dpERK (B and D) in synaptic boutons and tracheal cells in *repo-GAL4* and *repo-GAL4*-driven *UAS-decoy-mir*-*274* (A and B) and *mir-274*^*KO*^/*mir-274*^*KO*^; *repo-GAL4*/+ and *mir-274*^*KO*^/*mir-274*^*KO*^; *repo*>*mir-274* (C and D). (E-G) Bar graphs for quantifications of Sty (F and G) and dpERK (F and G) immunofluorescence intensities within synaptic boutons and tracheal cells and the numbers of synaptic boutons and tracheal branches (J). (E and F) Data was analyzed by Independent *t*-tests. (G) Data was analyzed by One-way ANOVA followed by Tukey post hoc tests.

Finally, we showed that disruption of the exosomal biogenesis, transportation and release by glial knockdowns of Rab11, Syx1A, TSG101 and Shrb also caused Sty upregulation and dpERK downregulation in synaptic boutons and tracheal cells (Figures S5A-H). Taken together, these results suggest that glia-derived miR-274 downregulates Sty and upregulates dpERK levels in synaptic boutons and tracheal cells.

### miR-274 modulates larval hypoxia response

The *Drosophila* trachea is a highly branched network with open ends and air-filled terminal branches. It functions in gas exchange similarly to mammalian circulatory systems (35, 36). Oxygen tension is important for inducing tracheal terminal branching (37). We postulated that miR-274 might play a physiological role in glia-modulated tracheal branching during hypoxia. To test this possibility, we assayed the hypoxia escape response for the *mir-274*^*KO*^ mutant (see Behavior assay in Materials and Methods) (38). When exposed to hypoxia (1% O_2_), only about 20% of control larvae (*w*^*1118*^ and *Canton S*) had responded strongly within 5 min by fleeing the food source, and this percentage increased to almost 40% by 10 min and to close to 50% by 15 min (Figure 7A). In contrast, almost 50% of *mir-274*^*KO*^ mutants exhibited a strong hypoxia response (fleeing the food source) by 5 min and about 60% by 10 and 15 min (Figure 7A). No significant differences were found when we assayed these three genotypes under normoxia as a control, with almost all larvae (> 95%) staying in the food source (Figure 7B). Moreover, we confirmed that the hypoxia-induced response is not due to differences in larval locomotion, as we observed comparable crawling lengths among *mir-274*^*KO*^ mutants and controls (Figure S6A). We performed additional control experiments to show that *mir-274*^*KO*^ larvae are indeed more responsive to lower oxygen levels. First, *mir-274*^*KO*^ mutants still exhibited a significantly different hypoxia response compared to control larvae in a ten-fold-diluted food source, suggesting that the enhanced exploratory behavior of mutant flies is not due to differences in evaluating nutrition (Figure S6B). Second, feeding motivation toward nutritious (yeast) or non-nutritious (grape juice) foods, as evaluated by counting mouth hook contractions, revealed almost identical results under both fed and starved conditions (Figure S6C). Finally, the difference we had observed between *mir-274*^*KO*^ and control larvae was still significant when we conducted the assay in 10% oxygen, suggesting that the mutant larvae exhibited hypersensitivity towards reduced oxygen levels (Figure S6D).

**Figure 7.**
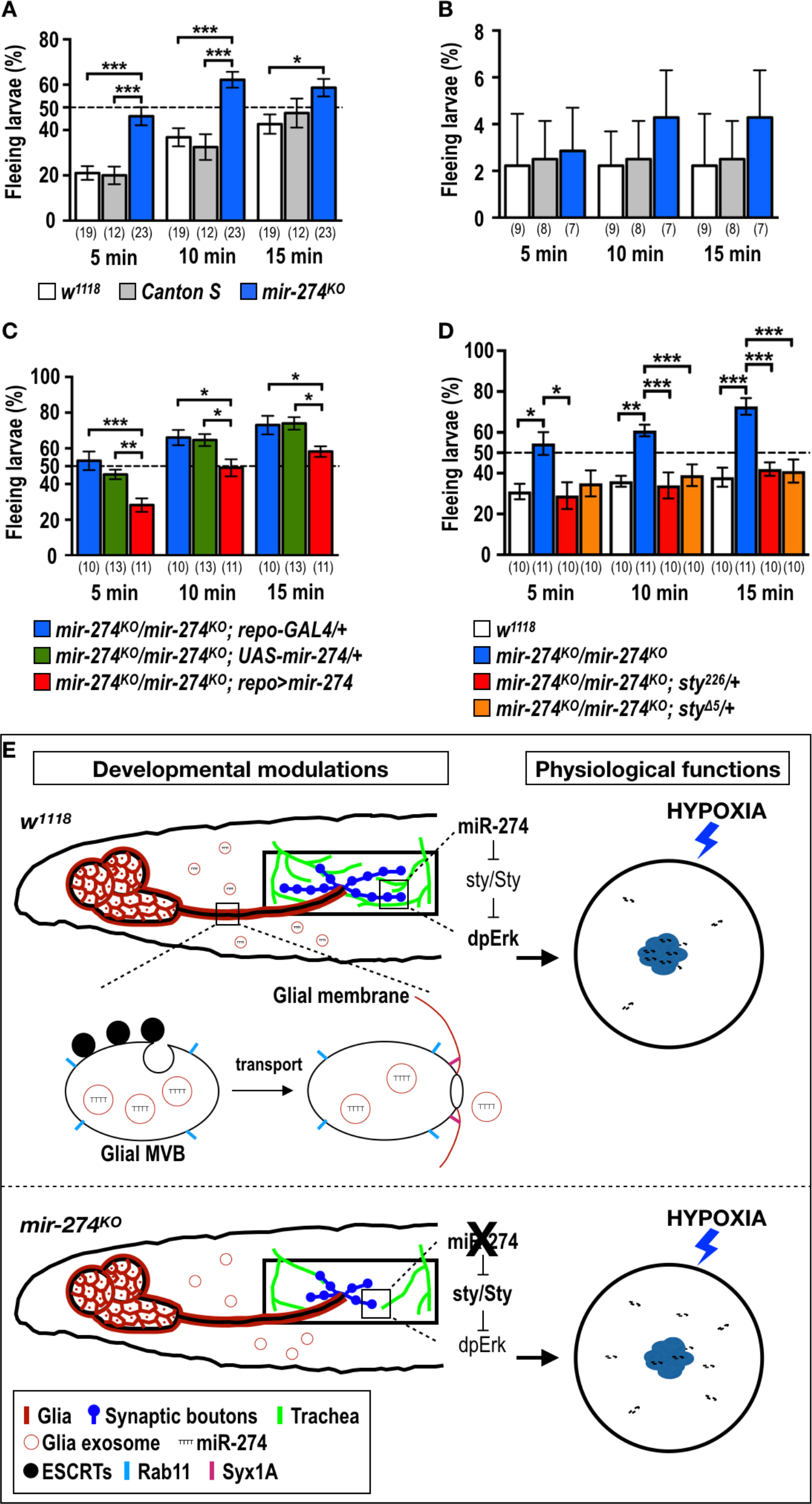
miR-274 modulates larval hypoxia responses. (A-D) Bar graphs showing quantifications of percentages of larvae fleeing a food source under conditions of 1% (A, C and D) or 21% (B) oxygen. (A and B) *w*^*1118*^, *Canton S*, and *mir-274*^*KO*^, (C) glia rescue (*mir-274*^*KO*^/*mir-274*^*KO*^; *repo*>*mir-274*) and two mutant controls (*mir-274*^*KO*^/*mir-274*^*KO*^; *repo-GAL4*/+ or *mir-274*^*KO*^/*mir-274*^*KO*^; *UAS-mir-274*/+), and (D) bi-allelic *sty* suppression (*mir-274*^*KO*^/*mir-274*^*KO*^; *sty*^*226*^/+ and *mir-274*^*KO*^/*mir-274*^*KO*^; *sty*^*Δ5*^/+). Data were analyzed by One-way ANOVA followed by Tukey post hoc tests. (E) Diagrams depicting how miR-274 is sorted into exosomes in MVBs through ESCRT-dependent mechanisms and then released extracellularly through Rab11- and Syx1A-dependent mechanisms. Secreted miR-274-containing exosomes downregulate Sty expression within targeting tissues, which in turn upregulates dpERK levels, thus promoting growth of synaptic boutons and tracheal branches. In the *miR-274* mutant, increased Sty expression downregulates dpERK levels and inhibits growth of target tissues. Mutant larvae with reduced tracheal branches are hypersensitive to hypoxic conditions.

We performed rescue experiments to examine whether glia-expressed miR-274 is required for normal larval hypoxia responses. Homozygous *mir-274*^*KO*^ mutants carrying both *repo-GAL4* and *UAS-mir-274* transgenes showed a significantly reduced hypoxia response compared to homozygous *mir-274*^*KO*^ mutants carrying either the *repo-GAL4* or *UAS-mir-274* transgenes (Figure 7C). As glial rescue larvae displayed a normal level of tracheal branching (Figures 1C and 1F), our results suggest that reduced tracheal branching might contribute to the altered hypoxia escape response. To examine whether miR-274 functions through Sty downregulation to affect this response, as for developmental regulation of tracheal branching, we performed genetic suppression of Sty by mutation. Indeed, we found that introducing a *sty* mutant allele in homozygous *mir-274*^*KO*^ mutants almost completely suppressed the enhanced hypoxia escape response (Figure 7D). Since tracheal branching was also restored by the *sty* mutant alleles in the absence of miR-274 (Figures 5E and 5H), this result supports that tracheal branching is linked to the hypoxia escape response.

## Discussion

Here, we report that glia-derived miR-274 non-cell-autonomously regulates synaptic and tracheal growth during development. miR-274 is expressed in glia and regulates Sty expression and MAPK signaling in synapses and trachea to control their growth. Exosome-borne miR-274 derived from glia was present in the circulatory hemolymph, which requires glial Rab11, Syx1A, and ESCRT complex components. We further show that miR-274 is present in synapses and trachea following specific glial expression of miR-274 in the *mir-274*^*KO*^ mutant. Glial expression of miR-274 also restored synapse and tracheal growth through Sty downregulation. We observed that *mir-274*^*KO*^ larvae with compromised tracheal branching were hypersensitive to hypoxia, which could be suppressed by reducing the *sty* gene dosage to reestablish the tracheal system. We propose that developmentally regulated neuron-trachea coupling via glial-secreted miR-274 is synchronized post-developmentally to physiological demands (Figure 7E).

### Circulating miR-274

Extracellular miRNAs detected in the blood serum and other body fluids are highly stable and resistant to RNase treatments, making them ideal signaling molecules for long-distance communication among tissues and organs (39–42). In *Drosophila*, miRNAs have been isolated from the circulating hemolymph (10), suggesting that *Drosophila* miRNAs are also carried by exosomes in the circulatory system to function systematically for tissue and organ interactions. Formation of cargo-carrying exosomes requires the ESCRT complexes, which facilitate membrane invagination and form the intraluminal vesicles within MVBs (27). Rab11 and Syx1A in neurons effectively block exosome secretion from presynaptic boutons by disrupting the exosome transportation and release machinery (30). We detected miR-274 in exosomal fractions of larval hemolymph (Figure 3D), even when we contrived to specifically express miR-274 solely in glia (Figure 3E). Furthermore, Rab11, Syx1A and partially the ESCRT components in glia were required to release miR-274 into the hemolymph (Figures 4A and S3A) and in target tissues (Figure 4B). These results support that glia-expressed miR-274 is carried by exosomes in the larval circulatory system.

Circulating exosomes carry diverse molecules including proteins and RNAs. Although some studies have suggested that extracellular miRNAs might be “cellular by-products” disposed of by apoptotic cells (42), our genetic data provide strong evidence of a non-cell-autonomous developmental role for glial miR-274. Though miR-274 may cell-autonomously regulate secreted factors in glia to execute its function indirectly, our findings support an active and direct role for miR-274 at target sites. In addition to confirming the presence of miR-274 at target tissues following glia-only expression (Figure 3G), we observed that expression of Sty that carries a miR-274 target site at its 3’UTR was suppressed in target sites upon glia-only miR-274 expression (Figure 6C). Circulating miR-274 in the hemolymph could potentially target multiple tissues given that we detected miR-274 as well as regulation of Sty and dpERK expression also in muscle cells (for example, see Figures 6C and 6D). The function of miR-274 in muscle cells awaits further study. Accordingly, miR-274 might have a systematic role in multiple tissues that could contribute to coordinating their developmental processes and post-developmental physiology.

### Glia specificity of miR-274 secretion

The non-cell-autonomous role of miR-274 appears to be highly cell-type specific. Although expression of pre-miR-274 is highly glia-enriched, perhaps accounting for the majority of specificity, other layers of regulation may confer this specificity. We have shown that miR-274 secretion is highly specific to glia, as glia-expressed miR-274 was detected in synaptic boutons, muscle and tracheal cells, whereas neuron- and trachea-expressed miR-274 was only detected in the respective expressing cells (Figures 3E, S2A and S2D). Interestingly, neuron-expressed miR-274 was also detected in muscle cells (Figure S2A), which might be transported by transverse exosomes crossing the synaptic cleft at NMJs (30). The Wnt/Wg signal is carried by Evenness interrupted (Evi)-positive exosomes from pre- to post-synapses (30). Developmental signals like Hedgehog are also transported over long distances in wing epithelia for cell fate induction (28). The *Drosophila* retrovirus-like Gag protein Arc1 (dArc1) binds to *darc1 mRNA* to be sorted into exosomes for transport across synaptic boutons (29). Interestingly, although presynaptic release of Wg/dArc1 and glial miR-274 shares a requirement for Rab11 and Syx1A, they may still exhibit substantially difference. Thus, multiple secretory exosomal pathways carry distinct cargos and function in different tissues of *Drosophila*.

Exosomes are formed through multiple pathways—such as ubiquitination-dependent and –independent or ESCRT-dependent and –independent pathways—that package different combinations of cargoes (2, 3). We did not detect miR-274 in hemolymph exosomes when it was specifically expressed in neurons or tracheal cells (Figure 3E), perhaps because neurons or tracheal cells do not generate miR-274-bearing exosomes. Furthermore, data from our cell-type-specific rescue experiments clearly indicate that only glial miR-274 expression in a *mir-274*^*KO*^ mutant had a profound rescue effect on synapse and tracheal growth (Figures 1C and 1F). In contrast, the minor rescue effects of neuronal and tracheal expressions of miR-274 were restricted to the expressing cells (Figures S2B, S2C, S2E and S2F).

It has been suggested that miRNAs are subjected to cell-type-specific modifications, including uridylation and adenylation that alter miRNA localization, stability or activity (43). Such modifications may further induce packaging of miRNAs into exosomes for secretion in glia. Glia-specific miR-274 release suggests another layer of regulation for exosome-mediated cell-cell communication. Cargo packaging and exosome formation pathways are distinct in different types of cells (2, 3). We observed differential effects of knocking down several ESCRT components in terms of regulating synapse and tracheal growth, which could reflect the existence of heterogeneous populations of exosomes (Figures S3B and S3C). Differential requirements for ESCRT components have also been observed for blocking Hh-borne exosomes from wing-disc epithelial cells (28), as well as in the presynaptic release of Evi-positive exosomes (30). Thus, it would seem that complex regulation of the biogenesis of distinct exosomal populations may underlie different exosome-mediated communications between specific pairs of source and target cells. Different miRNA species have been detected in exosomes isolated from various types of immune, cancer, adipose and glial cells (6–8). Our analysis of the non-cell-autonomous function of miR-274 serves as a foundation for further study of the cell- and tissue-specificity involved in exosome-mediated cell-cell communication.

### Glia-modulated growth of trachea branches and hypoxia responses

Similar to mammalian systems, *Drosophila* glia are linked to neurons and vascular systems in terms of their structure and function. In the larval *Drosophila* brain, trachea grow alongside glial processes toward the central neuropiles (22). In the peripheral nervous system of adult flies, glial processes are intertwined with synaptic bouton-bearing axonal terminals and tracheal terminal branches to form the functional complex (21). This coupling between tracheal and neuronal processes may ensure efficient oxygen supply to neurons for activity and homeostasis, which is similar to the coupling between the vascular and nervous systems in vertebrates. At NMJs, the gliotransmitters Wnt/Wg and TNF-α regulate synaptic plasticity (44, 45). Glia also function as macrophages, engulfing synaptic debris and shaping neurites after injury (46). Direct ablation of glia throughout development induces tracheal branching, suggesting that tracheal branching is restricted by glia (22). In this study, we further report the co-regulation of both tracheal and nervous systems by glial-derived miR-274, reinforcing the idea of glia-neurovascular coupling in *Drosophila*. We show that only blockage of miR-274 in glia, not other target tissues, significantly affected the growth of synaptic boutons and trachea.

Here, we chose Sty to investigate miR-274 targeting since Sty is a negative regulator of RTK/Ras/MAPK signaling and is involved in synaptic growth and tracheal branching (32–34). Synaptic bouton numbers are consistently reduced when *sty* is overexpressed in neurons (32), a phenotype recapitulated in the *mir-274*^*KO*^ mutant. Loss-of-function mutations in *sty* enhanced tracheal branching, with increases ranging from 20-70% depending on different allelic combinations (33). Thus, Sty is sensitive to modulation, such as by miR-274 as shown in our study. It would be of great interest to establish whether and how miR-274 is regulated throughout development and physiology. Nevertheless, direct (and therefore more effective) glial regulation is supported here by the fact that Sty levels in target sites were modulated by miR-274. Such direct modulation in target sites could ensure synchronized growth regulation for both synaptic boutons and tracheal branches.

Recently, miRNAs were also shown to be essential for physiological functions. In *Drosophila*, miR-iab4/iab8 is expressed in self-righting node neurons (SRNs) and is responsible for larval self-righting behavior. Lack of miR-iab4/iab8 or overexpressing the target gene *Ultrabithorax* in SRNs inhibits the ability of larvae to right themselves (47). Similarly, astrocyte-specific expression of miR-263b and miR-274 is essential for circadian locomotor activity rhythms (48). Here, we reveal that *mir-274*^*KO*^ mutants barely tolerated a sustained low oxygen environment. This behavioral defect was correlated with *mir-274*^*KO*^ mutants having fewer tracheal branches as it was rescued by *sty* genetic suppression to restore tracheal branching. Oxygen may be delivered in the body less efficiently by having fewer tracheal terminal branches, rendering mutants less tolerant to low oxygen levels. Thus, miR-274 seems to ensure a well-developed tracheal system (and perhaps also synaptic boutons), allowing larvae to tolerate hypoxia. Our study highlights a coordinating role for glia in regulating a coupled developmental and physiological process.

## Acknowledgements

We thank M. Krasnow, T. Kornberg, Y.-W. Chen, Bloomington Stock Center and Vienna *Drosophila* Resource Center for providing antibodies and fly stocks, the Genomics Core and Bioinformatics-Biology Service Core in Academia Sinica for performing NGS and respective analysis, and the Imaging Core in the Institute of Molecular Biology of Academia Sinica for supporting image processing by Imaris. This work was supported by grants from the Ministry of Science and Technology (MOST) and Academia Sinica to C.T.C and MOST-104-2633-B-029-001, MOST-105-2633-B-029-001 to Y.C.T. The authors declare no biomedical financial interests or potential conflicts of interest.

## Material and methods

### Fly stocks

All flies were reared at 25 °C under a 12:12 hr light:dark cycle. Third instar wandering larvae were used for experiments. Mutant flies including *mir-274*^*KO*^ (58904), *sty*^*Δ5*^ (6382) and *sty*^*226*^ (6383) were obtain from Bloomington *Drosophila* Stock Center (BDSC). *mir-274*^*6-3*^ was generated by Dr. Yu-Chen Tsai in this paper. The *mir-274*^*6-3*^ allele bears a mutation in the seeding motif. The GAL4/UAS system was used, with *repo-GAL4* (BDSC_7415), *elav-GAL4* (BDSC_458) and *btl-GAL4* (BDSC_8807) used to drive glia, neuron and tracheal expression of transgenes. UAS responders were: *UAS-mir-274* (BDSC_59895), *UAS-mCD8-GFP* (BDSC_5137) and *UAS-myr-RFP* (BDSC_7119). RNAi flies were obtained from BDSC or Vienna *Drosophila* Resource Center (VDSC) including: *UAS-TSG101*^*RNAi*^ (BDSC_35710), *UAS-shrb*^*RNAi*^ (BDSC_38305), *UAS-Vps28*^*RNAi*^ (VDRC_105124), *UAS-Chmp1*^*RNAi*^ (BDSC_28906), *UAS-ALiX*^*RNAi*^ (BDSC_33417), *UAS-Rab11*^*RNAi*^ (BDSC_27730) and *UAS-Syx1A*^*RNAi*^ (BDSC_25811). The lexA/lexAop system was used, *btl-lexA*-driven *lexAop-CD2-GFP* was used to present the pattern of tracheal branches, the gift of Dr. Kornberg (49). The Decoy-mir-274 construct was generated as described (50). In brief, decoy miR-274-binding sites were synthesized by PCR to assemble multiple copies of the Decoy-linker-mir-274 cassette. Three copies of the Decoy-linker-mir-274 cassette were integrated into the pUAST vector (*Drosophila* Genomics Resource Center) to generate the Decoy-mir-274 plasmid for producing transgenic lines.

### Immunostaining, microscopy and image processing

Dissected third instar larval fillets were fixed and stained as described (51). Primary antibodies used in this study were goat anti-HRP-conjugated TRITC (1:100; Jackson Lab), rabbit anti-HRP-conjugated 647 (1:100; Jackson Lab), chicken anti-GFP (1:500; Abcam), mouse anti-Dlg (1:100; Developmental Studies Hybridoma Bank (DSHB)), mouse anti-activated MAP kinase (diphosphorylated ERK-1&2, 1:20; Sigma-Aldrich), and rabbit anti-Sty (1:50; gift of Dr. Krasnow (33)). Muscles were visualized by TRITC-conjugated phalloidin (1:2000; Sigma-Aldrich). Tracheal nuclei were visualized by DAPI (1:5000; Sigma-Aldrich). Goat anti-chicken (Invitrogen) or mouse Alexa Fluor 488-(Jackson Lab), goat anti-rabbit Cy5-(Jackson Lab) and goat anti-mouse Cy5-conjugated (Jackson Lab) secondary antibodies were used at 1:500 dilutions. Z-stack confocal images were obtained by Zeiss LSM510 or LSM 710 microscopy and processed by Imaris 8.4.1 (Bitplane).

Fiji (https://fiji.sc/) was used to quantify the signal intensities of Sty and dpERK in synaptic boutons, tracheal branches or cell nuclei. In brief, fluorescent integrated densities of Sty or dpERK were measured within the regions of interest (ROI), i.e., HRP-labeled synaptic boutons, GFP-labeled tracheal branches, or DAPI-labeled tracheal nuclei. The integrated densities were normalized to background readings of respective images.

For electron microscopy imaging, ultrastructures of dissected third instar larvae were processed as described to reveal glia-neuron-trachea organization (51).

### Fluorescent *in situ* hybridization (FISH)

Wandering larvae were dissected to perform EDC-fixed FISH as described (52–54). To detect precursor miR-274, we designed a 5’-biotin-labeled probe complementary to the loop sequence. To design a probe for mature miR-274 and a scramble probe, we used a 5’-DIG-labeled miRCURY LNA probe complementary (or scrambled) to the stem sequence (Exiqon). After hybridization, samples were blocked in Western blocking (WB):PBT (1:1) solution (Sigma-Aldrich) for 1 hr and incubated overnight with sheep anti-DIG-POD primary antibody (1:1000, for the DIG-labeled probe; Roche) or mouse anti-biotin (1:1000, for the biotin-labeled probe; Abcam) in the WB:PBT solution at 4 °C. Samples with the biotinylated probe were incubated for 2 hr in biotinylated donkey anti-mouse secondary antibody (1:500; Jackson Lab) in WB:PBT solution, washed three times in PBS for 10 min, and then incubated in HRP-labeled avidin-biotin complex (ABC; Vector Laboratories) solution for 1 hr. After washing three times in PBT, each time for 10 min, all samples were incubated in TSA Plus Cyanine5 Evaluation kit (1:50; PerkinElmer) for signal amplification at room temperature for 1 hr with protection from light. Finally, the samples were washed six times in PBT, each time for 5 min. To double-stain proteins of interest, we incubated the samples in 0.1% hydrogen peroxide in PBT for 10 min to eliminate enzymatic activity in the TSA Plus Cyanine5 Evaluation kit, followed by standard immunostaining procedures (51).

### Exosome isolation and Western blot

Exosome fractions were isolated from *Drosophila* S2 cell culture (2 × 10^6^ cells/ml) medium and the hemolymph of 50 to 100 larvae. In brief, the S2 cell culture medium or larval extracellular fluid was centrifuged at 300 *g* for 5 min, the supernatant was next spun at 2,000 *g* for 10 min and then at 10,000 *g* for 30 min to remove large cell debris. Exosomes were then collected following the manufacturer’s instructions for the ExoQuick™kit (System Biosciences). For Western blot analysis, we homogenized exosome fractions in RIPA lysis buffer with protease inhibitor cocktails. Total protein (40 µg) was loaded in each lane of 4-12% Nupage Bis-Tris gels. We used mouse anti-TSG101 (1:1000; Abcam), mouse anti-Syntaxin (1:1000; DSHB), mouse anti-Rab11 (1:1000; BD Biosciences) and horseradish peroxidase conjugated anti-mouse IgG (1:5000; Jackson Lab) for immunostaining.

### RT-PCR and real-time PCR

Total RNA was extracted form larvae or exosomal fractions using the miRNeasy Mini Kit (Qiagen) according to the manufacturer’s instructions. SuperScript™ IV VILO™ Master Mix (ThermoFisher Scientific) was used according to the manufacturer’s instructions to detect mRNAs. To detect miR-274, a SuperScript™ III Reverse Transcriptase (ThermoFisher Scientific) protocol was performed according to the manufacturer’s instructions with the stem-loop RT primer that contains six nucleotides complementary to mature miR-274 3’ sequences. For each reverse transcription reaction, 2 µg RNA was used. Phusion Green High-Fidelity DNA Polymerase (ThermoFisher Scientific) was used to perform RT-PCR with specific primer sets for miR-274 and *Ephrin* (Figures 4A, 5A and S4). To measure *sty* mRNA levels, we performed real-time PCR in 96-well plates using the LightCycler^®^ 480 system. All mRNA transcripts were normalized to *Rpl19* transcript levels. To measure miR-274 levels (Figures 4B, 5B and S5C), we conducted absolute qPCR by generating 8-point standard curves with miR-274 oligonucleotides in the range of 1 μM to 10^−7^ μM. Amounts of miR-274 were normalized to each respective *GAL4* control group and are shown as “fold change”. All primers were listed in Table 1.

**Table 1.**
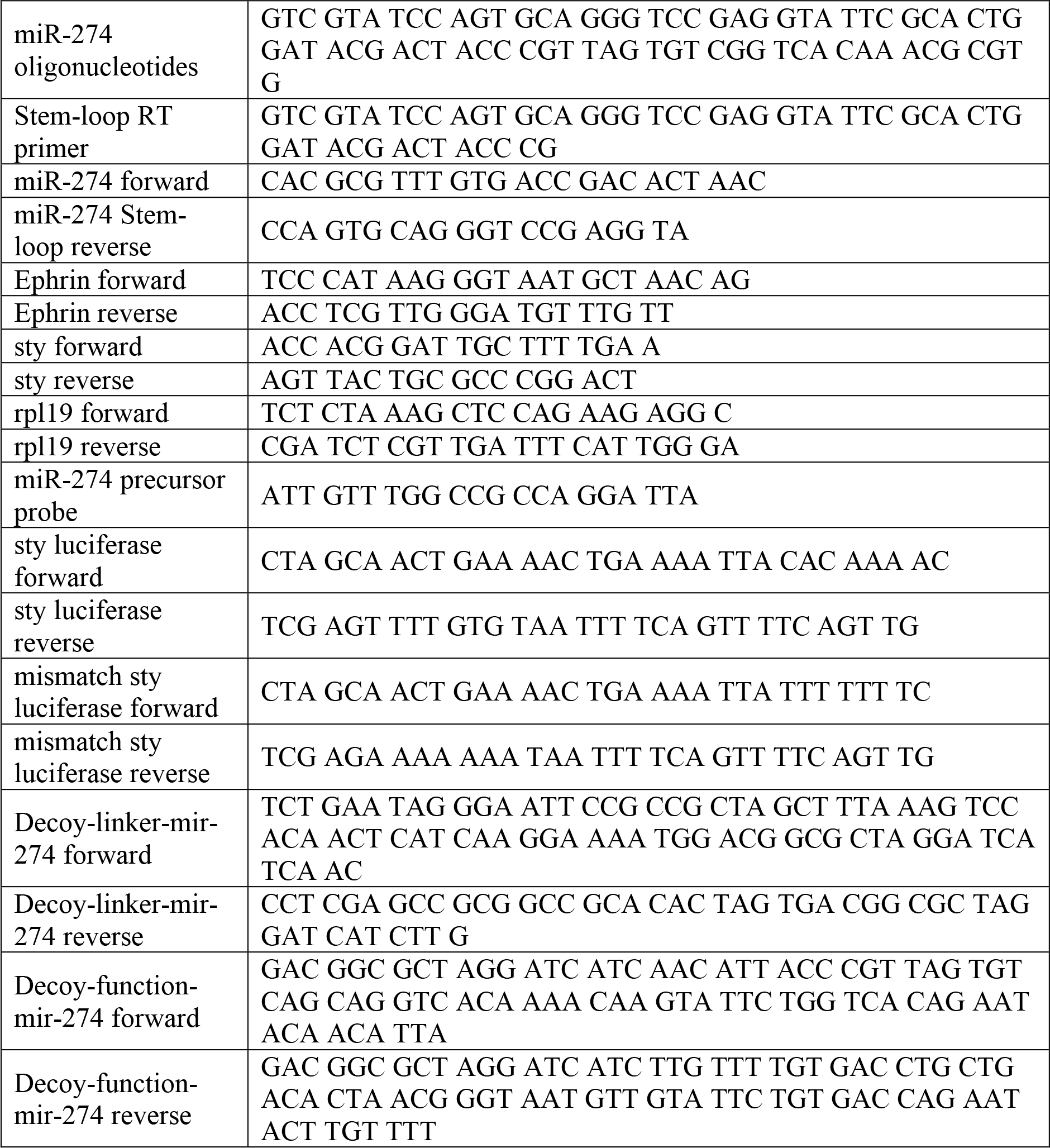
List of oligonucleotides.

### RNA sequencing

Total RNA was extracted from wandering female larvae of *w*^*1118*^ and *mir-274*^*KO*^ using a miRNeasy Mini Kit. Libraries were prepared with a TruSeq Stranded mRNA LT Sample Prep Kit for Illumina sequencing. We selected genes in the *mir-274*^*KO*^ mutant with expression levels 1.5-fold those of *w*^*1118*^.

### Luciferase construct and assay

Oligonucleotides targeting predicted sequences of *miR-274* and the mismatch control at the *sty* 3’-UTR were synthesized and inserted into the NheI/XhoI multiple cloning site of pmirGLO Dual-Luciferase miRNA target expression vector (Promega). Three plasmid constructs were generated: (1) without 3’-UTR sequences; (2) with wild-type 3’-UTR sequences; and (3) with mismatched 3’-UTR sequences. These plasmids (1 µg each) were transfected by the TransIT system (Mirus) into 2×10^6^ S2 cells that were plated in 6-well plates one day before analysis. Dual-luciferase reporter assays (Promega)were performed 48 hr after transfection according to the manufacturer’s instructions and measured by an EnSpire^®^ Multimode Plate Reader.

### Behavioral assay

Early- to mid-third instar larvae (AEL 72-96 hr) were used for the following behavioral assays.

#### Larval crawling assay

Individual larva was transferred to a 15 cm petri dish with 2% agarose for 1 min habituation. Activity was then video-recorded for 1 min using a Canon N90 camera and analyzed by Fiji with the wrMTrck plugin (55).

#### Feeding motivation assay

Larvae that were either well fed or food-deprived for 2 hr were transferred to a nutritious yeast or non-nutritious grape juice plate. Mouth-hook contractions of individual larvae were counted under dissecting microscopes for 1 min.

#### Hypoxia escape response

Hypoxia escape behavior was assessed as described (38), but with the following modifications. We used 10 larvae that had burrowed for 10 min into yeast paste applied to a grape juice agar plate. The assay was performed in a plexiglass chamber (20 × 10 × 10 cm) with O_2_ levels of 21%, 10% or 1% regulated by N_2_ infusion, which were monitored and controlled by a Proox 110 (BioSpherix) compact oxygen controller. Activity was video-recorded for 20 min using a Canon N90 camera and the number of larvae that moved beyond the yeast paste was counted at 5, 10, and 15 min.

### Statistical analysis

Graphpad Prism v6 (Graphpad) was used to perform statistical analyses. All data are expressed as mean ± SEM. The threshold for statistical significance was set at *p* < 0.05, with * for *p* < 0.05, ** for *p* < 0.01, *** for *p* < 0.001. Two-way ANOVA was used to analyze feeding motivation assay data. One-way ANOVA followed by Tukey post hoc tests were used to analyze data for three or more genotypes. Independent *t*-tests were used where appropriate.

---

**Figure S1.**
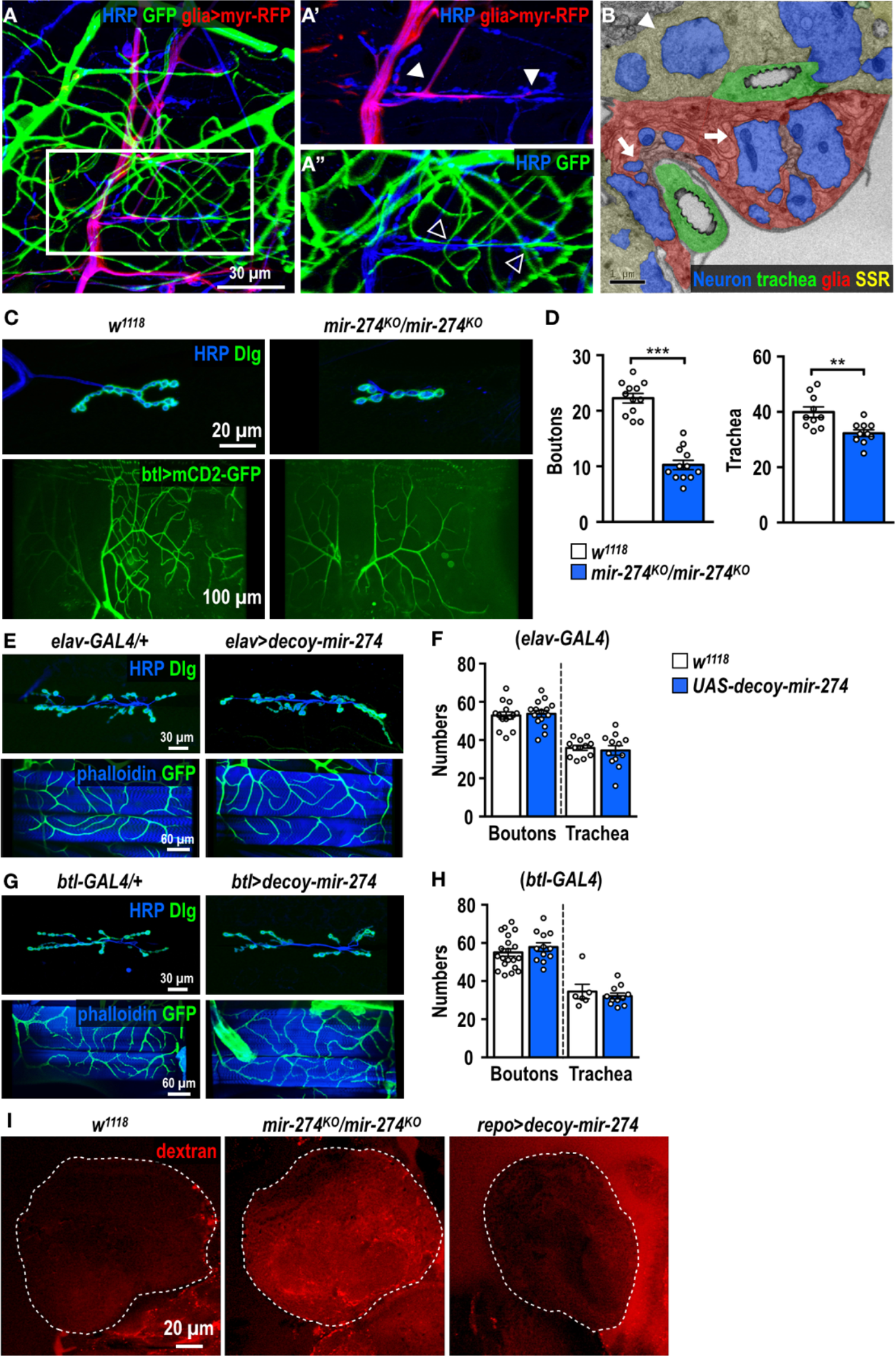
Growth modulation of synaptic boutons and tracheal branches by miR-274. (A-A”) Confocal images showing the glia-neuron-trachea coupling at NMJs of muscle 6/7 for screening miRNA mutants. Larvae were dissected to view glia labeled by *repo-GAL4*>*UAS-myr-RFP*, trachea labeled by *btl-lexA*>*lexAop-CD2-GFP*, and nerves stained by HRP. Boxed area in (A) is amplified to show that (A’) glial processes wrap around motor axons but not synaptic boutons (white arrowheads), and (A”) tracheal branches localize close to synaptic boutons (empty arrowheads). (B) Transmission election microscopy micrograph showing that glial processes directly enwrap axons, as indicated by arrows. Synaptic boutons (arrowhead) are recognized by the presence of synaptic vesicles, and surrounded by SSR. Coupling of tracheal branches to synaptic boutons, glial processes and axons is evident. (C) Images of synaptic boutons immunostained for presynaptic HRP and postsynaptic Dlg at NMJs of muscle 4 (top panels), and dorsal tracheal branches revealed by *btl-lexA*>*lexAop-CD2-GFP* (bottom panels) in *w*^*1118*^ and *mir-274*^*KO*^/*mir-274*^*KO*^. (E and G) Images of synaptic boutons and tracheal branches for *elav-GAL4* (E) and *btl-GAL4* (G) controls (crossed to *w*^*1118*^, left panels) or for miR-274 depletion (crossed to *UAS-decoy-mir-274*, right panels). (D, F and H) Bar graphs for quantifications of synaptic boutons and tracheal branches. Data was analyzed by Independent *t*-tests. (I) Leakage of dextran dye (24) into the larval brain (dotted line) was observed in *mir-274*^*KO*^, but not in *w*^*1118*^ or glia-driven *UAS-decoy-mir-274*. All scale bars are shown at bottom right.

**Figure S2.**
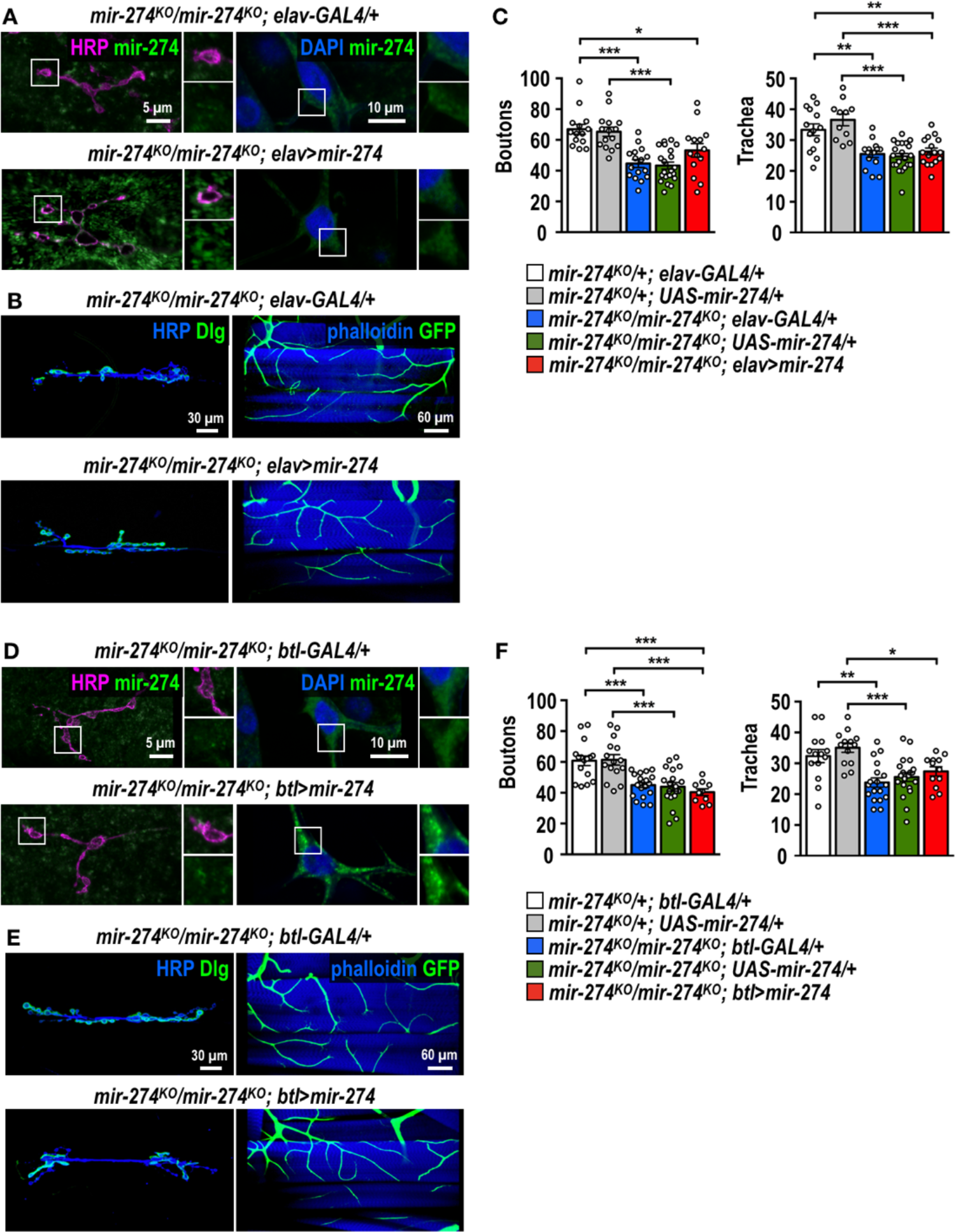
Restricted effects of neuronal and tracheal miR-274. (A and D) Confocal images showing background miR-274 signals generated by the mature probe in (A) *mir-274*^*KO*^/*mir-274*^*KO*^; *elav-GAL4* and (D) *mir-274*^*KO*^/*mir-274*^*KO*^; *btl-GAL4*. However, miR-274 FISH signals were detected in synaptic boutons of *mir-274*^*KO*^/*mir-274*^*KO*^; *elav*>*mir-274* (A) and tracheal cells of *mir-274*^*KO*^/*mir-274*^*KO*^; *btl*>*mir-274* (D). (B and E) Confocal images of synaptic boutons and tracheal branches upon neuronal (B) and tracheal (E) expression of miR-274, and in respective GAL4 controls. (C and F) Bar graphs for quantifications of synaptic bouton and tracheal branch numbers. Data was analyzed by One-way ANOVA followed by Tukey post hoc tests.

**Figure S3.**
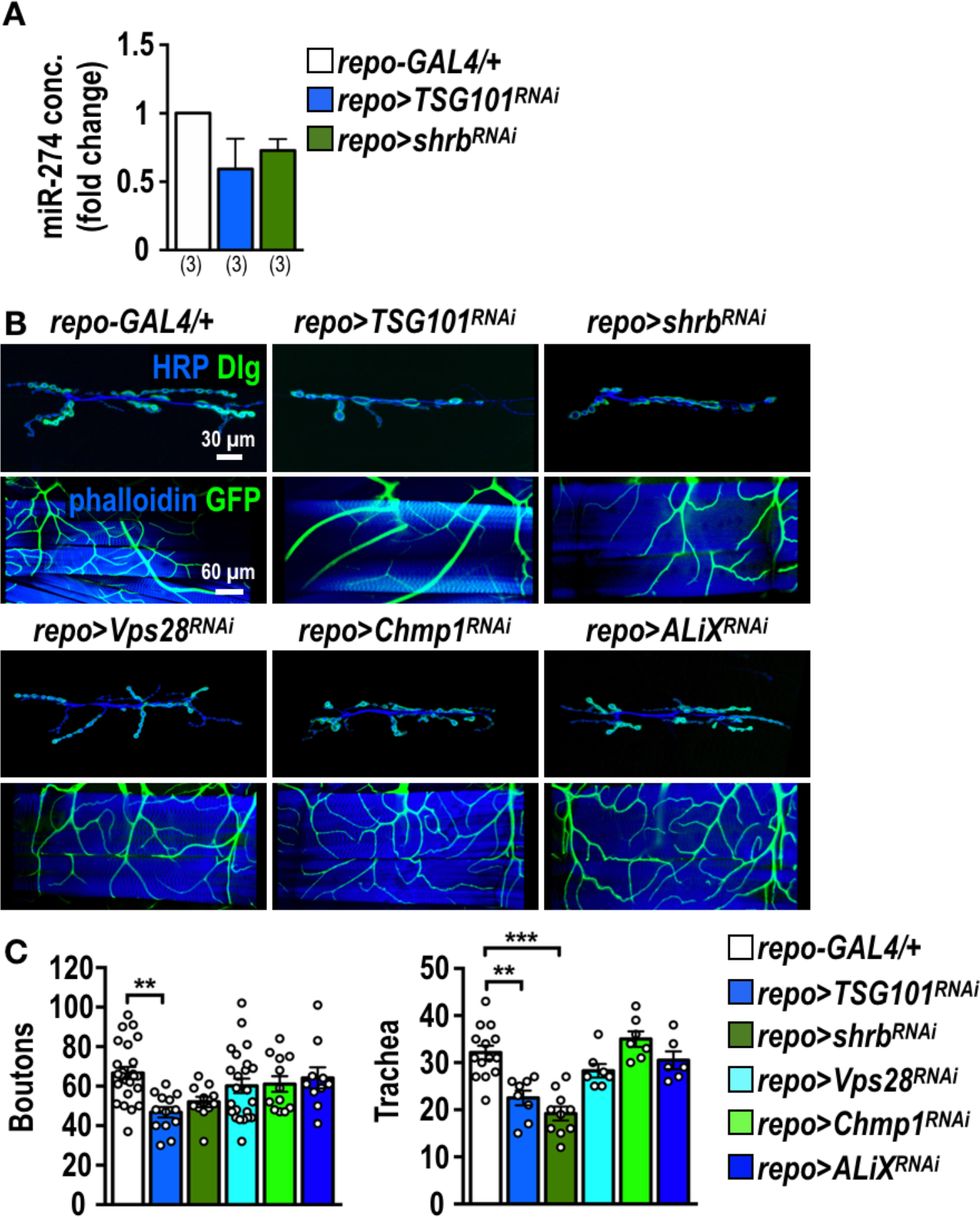
Secretion of exosomal miR-274 requires ESCRT in glia. (A) Absolute qPCR was performed to detect exosomal miR-274 in hemolymphs prepared from *repo-GAL4* and knockdowns of TSG101 and Shrb. (B) Confocal images of synaptic boutons and tracheal branches in *repo-GAL4*, *repo*>*TSG101*^*RNAi*^, *repo*>*shrb*^*RNiA*^, *repo*>*Vps28*^*RNAi*^, *repo*>*Chmp1*^*RNAi*^ and *repo*>*ALiX*^*RNAi*^. (C) Bar graph for quantification of synaptic boutons and tracheal branches. Data were analyzed by One-way ANOVA followed by Tukey post hoc tests.

**Figure S4.**
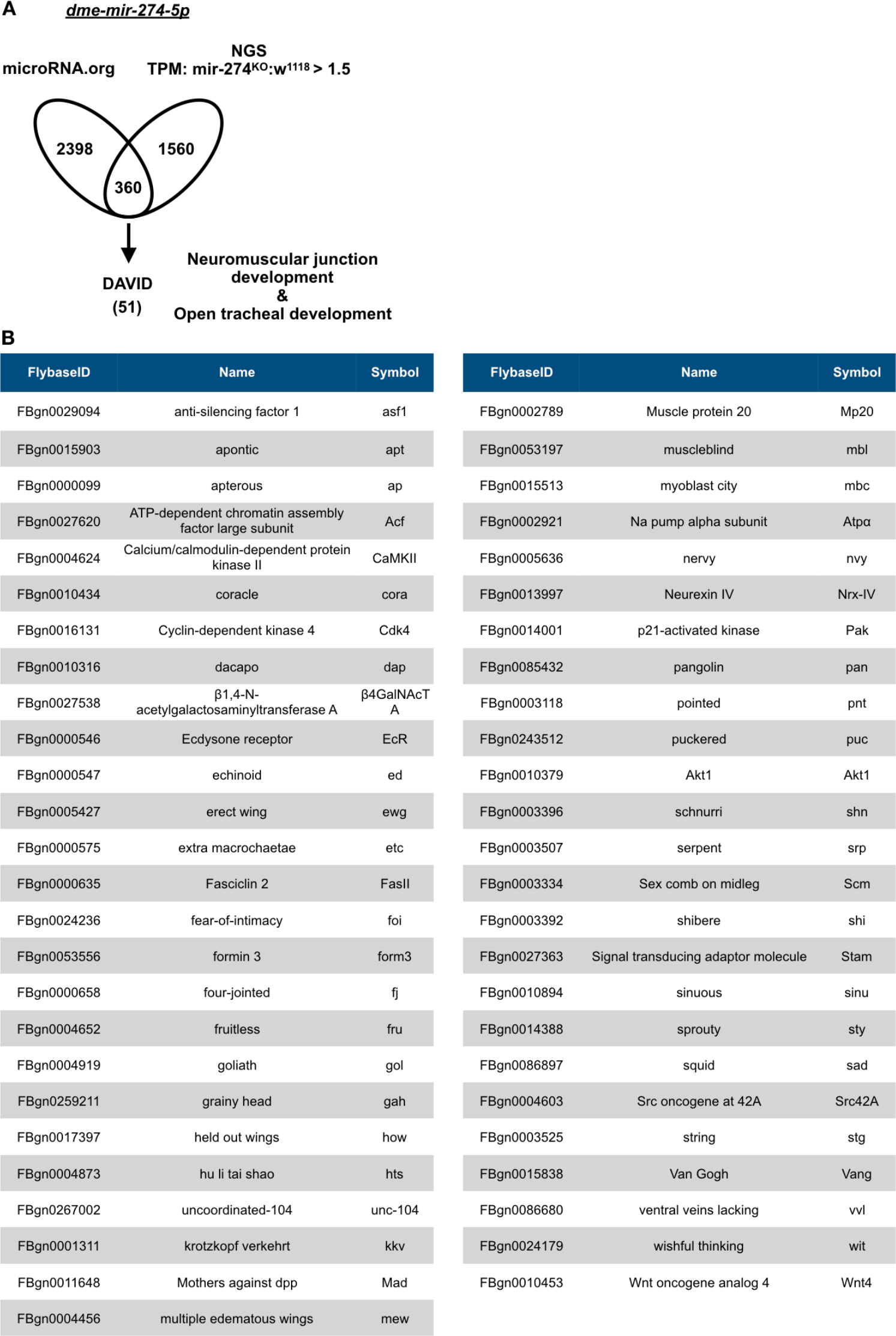
miR-274 target candidates. (A) Experimental approaches for miR-274 target selection. In one approach, potential miR-274 target genes were predicted and annotated by miRanda from www.microRNA.org. In another approach, genes upregulated in *mir-274*^*KO*^ larvae (>1.5-fold relative to *w*^*1118*^) were selected. Overlapping candidates were further filtered by DAVID Bioinformatics Resources 6.7 (https://david-d.ncifcrf.gov) for functions in NMJ and tracheal development. (B) The list of 51 candidate genes.

**Figure S5.**
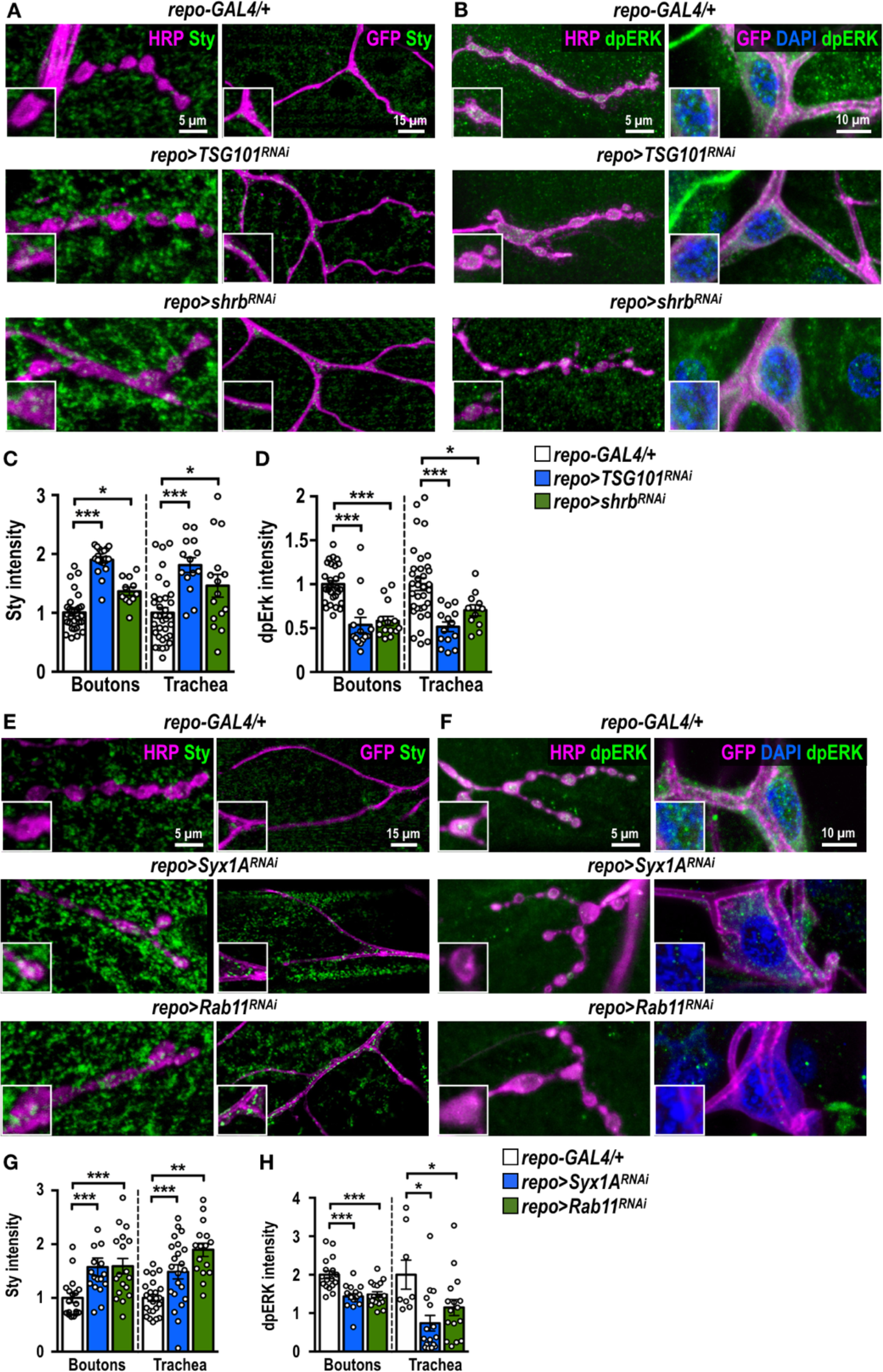
Exosomal secretion of miR-274 requires ESCRT conponents, Rabb11 and Syx1A to modulate synaptic and tracheal growth through Sty. (A, B, E and F) Confocal images showing immunostaining of Sty (A and E) or dpERK (B and F) in synaptic boutons and tracheal branches in *repo-GAL4*, *repo>TSG101*^*RNAi*^ and *repo*>*shrb*^*RNAi*^ (A and B) and repo>*Syx1A*^*RNAi*^ and *repo*>*Rab11*^*RNAi*^ (E and F). (C, D, G and H) Bar graphs for quantifications of Sty (C and G) and dpERK (D and H) immunofluorescence intensities within synaptic boutons and tracheal cells. Data were analyzed by One-way ANOVA followed by Tukey post hoc tests.

**Figure S6.**
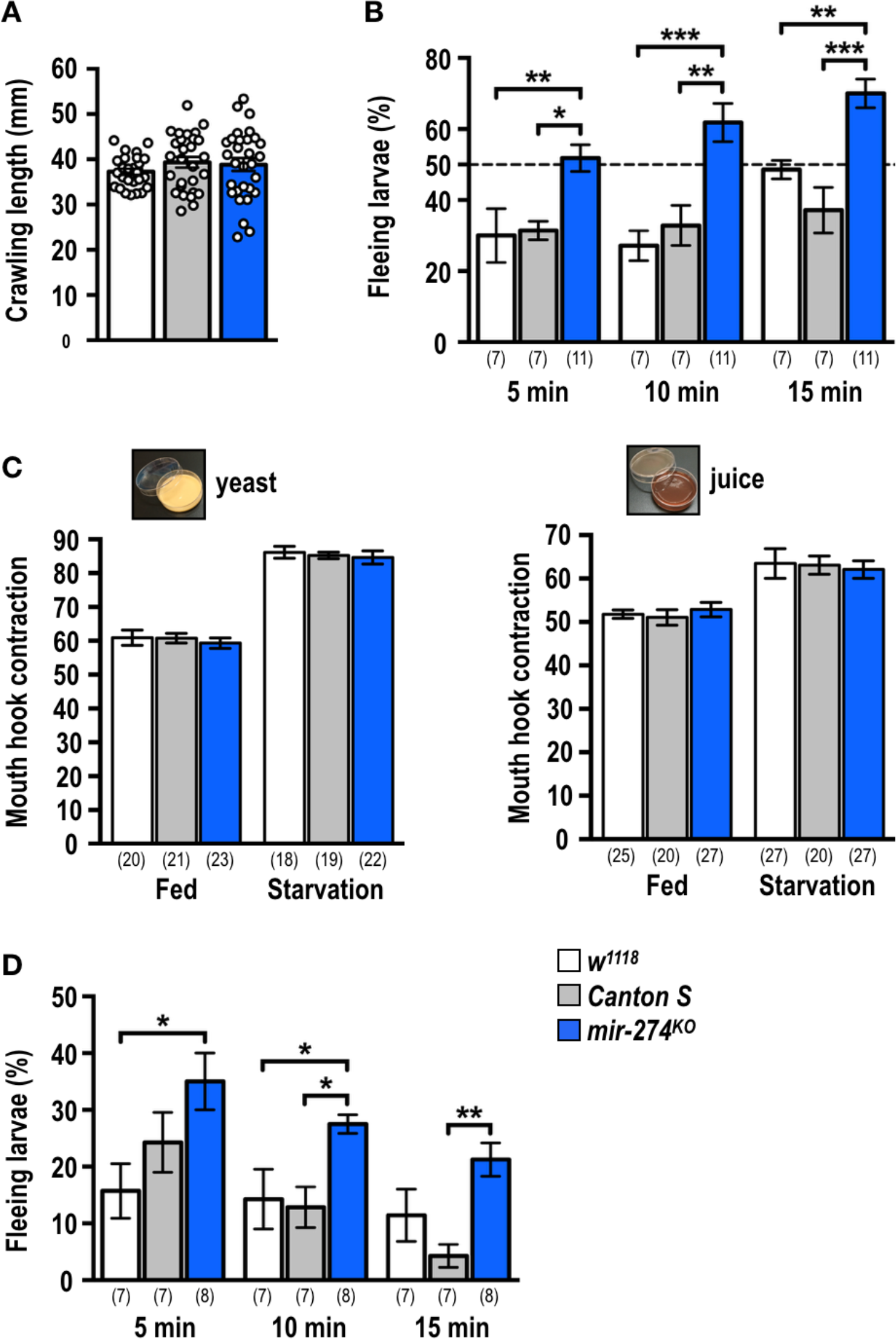
Responses of miR-274 mutant larvae to different environmental cues. (A) Bar graph showing comparable crawling lengths in 1 min among *w*^*1118*^ (white bar), *Canton S* (grey bar) and *mir-274*^*KO*^ (blue bar) larvae. (B) Bar graph showing quantification of larval fleeing behavior in response to 1% O_2_ with 1% nutrition. (C) Bar graph showing no significant differences among three genotypes in terms of the numbers of mouth hook contractions when larvae were fed or starved prior to being placed in yeast or grape juice plates. (D) Bar graph showing quantifications for the percentages of fleeing larvae under conditions of 10% O_2_. (A, B and D) Data were analyzed by One-way ANOVA followed by Tukey post hoc tests. (C) Data were analyzed by Two-way ANOVA.

## References

1. Hessvik NP & Llorente A (2018) Current knowledge on exosome biogenesis and release. Cell Mol Life Sci 75(2):193–208.

2. McGough IJ & Vincent JP (2016) Exosomes in developmental signalling. Development 143(14):2482–2493.

3. van Niel G, D’Angelo G, & Raposo G (2018) Shedding light on the cell biology of extracellular vesicles. Nature reviews. Molecular cell biology 19(4):213–228.

4. Thery C (2011) Exosomes: secreted vesicles and intercellular communications. F1000 Biol Rep 3:15.

5. Ha M & Kim VN (2014) Regulation of microRNA biogenesis. Nature reviews. Molecular cell biology 15(8):509–524.

6. Mittelbrunn M, et al. (2011) Unidirectional transfer of microRNA-loaded exosomes from T cells to antigen-presenting cells. Nature communications 2:282.

7. van der Vos KE, et al. (2016) Directly visualized glioblastoma-derived extracellular vesicles transfer RNA to microglia/macrophages in the brain. Neuro Oncol 18(1):58–69.

8. Thomou T, et al. (2017) Adipose-derived circulating miRNAs regulate gene expression in other tissues. Nature 542(7642):450–455.

9. Parrish JZ, Xu P, Kim CC, Jan LY, & Jan YN (2009) The microRNA bantam functions in epithelial cells to regulate scaling growth of dendrite arbors in drosophila sensory neurons. Neuron 63(6):788–802.

10. Dhahbi JM, et al. (2016) MicroRNAs Circulate in the Hemolymph of Drosophila and Accumulate Relative to Tissue microRNAs in an Age-Dependent Manner. Genomics Insights 9:29–39.

11. Tam SJ & Watts RJ (2010) Connecting vascular and nervous system development: angiogenesis and the blood-brain barrier. Annual review of neuroscience 33:379–408.

12. Carmeliet P & Tessier-Lavigne M (2005) Common mechanisms of nerve and blood vessel wiring. Nature 436(7048):193–200.

13. James JM, Gewolb C, & Bautch VL (2009) Neurovascular development uses VEGF-A signaling to regulate blood vessel ingression into the neural tube. Development 136(5):833–841.

14. Schwarz Q, et al. (2004) Vascular endothelial growth factor controls neuronal migration and cooperates with Sema3A to pattern distinct compartments of the facial nerve. Genes & development 18(22):2822–2834.

15. Vazquez AL, Masamoto K, Fukuda M, & Kim SG (2010) Cerebral oxygen delivery and consumption during evoked neural activity. Frontiers in neuroenergetics 2:11.

16. Florian C, Vecsey CG, Halassa MM, Haydon PG, & Abel T (2011) Astrocyte-derived adenosine and A1 receptor activity contribute to sleep loss-induced deficits in hippocampal synaptic plasticity and memory in mice. The Journal of neuroscience: the official journal of the Society for Neuroscience 31(19):6956–6962.

17. Henneberger C, Papouin T, Oliet SH, & Rusakov DA (2010) Long-term potentiation depends on release of D-serine from astrocytes. Nature 463(7278):232–236.

18. Mulligan SJ & MacVicar BA (2004) Calcium transients in astrocyte endfeet cause cerebrovascular constrictions. Nature 431(7005):195–199.

19. Nedergaard M, Ransom B, & Goldman SA (2003) New roles for astrocytes: redefining the functional architecture of the brain. Trends in neurosciences 26(10):523–530.

20. Newman EA (2003) New roles for astrocytes: regulation of synaptic transmission. Trends in neurosciences 26(10):536–542.

21. Danjo R, Kawasaki F, & Ordway RW (2011) A tripartite synapse model in Drosophila. PloS one 6(2):e17131.

22. Pereanu W, Spindler S, Cruz L, & Hartenstein V (2007) Tracheal development in the Drosophila brain is constrained by glial cells. Developmental biology 302(1):169–180.

23. Stork T, et al. (2008) Organization and function of the blood-brain barrier in Drosophila. The Journal of neuroscience: the official journal of the Society for Neuroscience 28(3):587–597.

24. Brink DL, Gilbert M, Xie X, Petley-Ragan L, & Auld VJ (2012) Glial processes at the Drosophila larval neuromuscular junction match synaptic growth. PloS one 7(5):e37876.

25. Menon KP, Carrillo RA, & Zinn K (2013) Development and plasticity of the Drosophila larval neuromuscular junction. Wiley interdisciplinary reviews. Developmental biology 2(5):647–670.

26. Chen YW, et al. (2014) Systematic study of Drosophila microRNA functions using a collection of targeted knockout mutations. Developmental cell 31(6):784–800.

27. Henne WM, Buchkovich NJ, & Emr SD (2011) The ESCRT pathway. Developmental cell 21(1):77–91.

28. Gradilla AC, et al. (2014) Exosomes as Hedgehog carriers in cytoneme-mediated transport and secretion. Nature communications 5:5649.

29. Ashley J, et al. (2018) Retrovirus-like Gag Protein Arc1 Binds RNA and Traffics across Synaptic Boutons. Cell 172(1-2):262–274 e211.

30. Koles K, et al. (2012) Mechanism of evenness interrupted (Evi)-exosome release at synaptic boutons. The Journal of biological chemistry 287(20):16820–16834.

31. Bergmann A, Agapite J, McCall K, & Steller H (1998) The Drosophila gene hid is a direct molecular target of Ras-dependent survival signaling. Cell 95(3):331–341.

32. Franciscovich AL, Mortimer AD, Freeman AA, Gu J, & Sanyal S (2008) Overexpression screen in Drosophila identifies neuronal roles of GSK-3 beta/shaggy as a regulator of AP-1-dependent developmental plasticity. Genetics 180(4):2057–2071.

33. Hacohen N, Kramer S, Sutherland D, Hiromi Y, & Krasnow MA (1998) sprouty encodes a novel antagonist of FGF signaling that patterns apical branching of the Drosophila airways. Cell 92(2):253–263.

34. Kramer S, Okabe M, Hacohen N, Krasnow MA, & Hiromi Y (1999) Sprouty: a common antagonist of FGF and EGF signaling pathways in Drosophila. Development 126(11):2515–2525.

35. Affolter M, et al. (2003) Tube or not tube: remodeling epithelial tissues by branching morphogenesis. Developmental cell 4(1):11–18.

36. Uv A, Cantera R, & Samakovlis C (2003) Drosophila tracheal morphogenesis: intricate cellular solutions to basic plumbing problems. Trends in cell biology 13(6):301–309.

37. Jarecki J, Johnson E, & Krasnow MA (1999) Oxygen regulation of airway branching in Drosophila is mediated by branchless FGF. Cell 99(2):211–220.

38. Wingrove JA & O’Farrell PH (1999) Nitric oxide contributes to behavioral, cellular, and developmental responses to low oxygen in Drosophila. Cell 98(1):105–114.

39. Chen X, et al. (2008) Characterization of microRNAs in serum: a novel class of biomarkers for diagnosis of cancer and other diseases. Cell Res 18(10):997–1006.

40. Chim SS, et al. (2008) Detection and characterization of placental microRNAs in maternal plasma. Clin Chem 54(3):482–490.

41. Mitchell PS, et al. (2008) Circulating microRNAs as stable blood-based markers for cancer detection. Proceedings of the National Academy of Sciences of the United States of America 105(30):10513–10518.

42. Turchinovich A, Tonevitsky AG, & Burwinkel B (2016) Extracellular miRNA: A Collision of Two Paradigms. Trends Biochem Sci 41(10):883–892.

43. Kim YK, Heo I, & Kim VN (2010) Modifications of small RNAs and their associated proteins. Cell 143(5):703–709.

44. Kerr KS, et al. (2014) Glial wingless/Wnt regulates glutamate receptor clustering and synaptic physiology at the Drosophila neuromuscular junction. The Journal of neuroscience: the official journal of the Society for Neuroscience 34(8):2910–2920.

45. Keller LC, et al. (2011) Glial-derived prodegenerative signaling in the Drosophila neuromuscular system. Neuron 72(5):760–775.

46. MacDonald JM, et al. (2006) The Drosophila cell corpse engulfment receptor Draper mediates glial clearance of severed axons. Neuron 50(6):869–881.

47. Picao-Osorio J, Johnston J, Landgraf M, Berni J, & Alonso CR (2015) MicroRNA-encoded behavior in Drosophila. Science 350(6262):815–820.

48. You S, Fulga TA, Van Vactor D, & Jackson FR (2018) Regulation of Circadian Behavior by Astroglial MicroRNAs in Drosophila. Genetics 208(3):1195–1207.

49. Roy S, Huang H, Liu S, & Kornberg TB (2014) Cytoneme-mediated contact-dependent transport of the Drosophila decapentaplegic signaling protein. Science 343(6173):1244624.

50. Haraguchi T, Ozaki Y, & Iba H (2009) Vectors expressing efficient RNA decoys achieve the long-term suppression of specific microRNA activity in mammalian cells. Nucleic Acids Res. 37(6):e43.

51. Wang CH, et al. (2017) USP5/Leon deubiquitinase confines postsynaptic growth by maintaining ubiquitin homeostasis through Ubiquilin. Elife 6.

52. Kerstens HM, Poddighe PJ, & Hanselaar AG (1995) A novel in situ hybridization signal amplification method based on the deposition of biotinylated tyramine. J. Histochem. Cytochem. 43(4):347–352.

53. Zimmerman SG, Peters NC, Altaras AE, & Berg CA (2013) Optimized RNA ISH, RNA FISH and protein-RNA double labeling (IF/FISH) in Drosophila ovaries. Nature protocols 8(11):2158–2179.

54. Pena JTG, et al. (2009) miRNA in situ hybridization in formaldehyde and EDC-fixed tissues. Nat. Meth. 6(2):139–141.

55. Nussbaum-Krammer CI, Neto MF, Brielmann RM, Pedersen JS, & Morimoto RI (2015) Investigating the spreading and toxicity of prion-like proteins using the metazoan model organism C. elegans. Journal of visualized experiments: JoVE (95):52321.

